# rCRUX: A Rapid and Versatile Tool for Generating Metabarcoding Reference libraries in R

**DOI:** 10.1101/2023.05.31.543005

**Authors:** Emily E. Curd, Luna Gal, Ramon Gallego, Shaun Nielsen, Zachary Gold

## Abstract

Key to making accurate taxonomic assignments are curated, comprehensive reference barcode databases. However, the generation and curation of such databases has remained challenging given the large and continuously growing volumes of DNA sequence data and novel reference barcode targets. Monitoring and research applications require a greater diversity of specialized gene regions and targeted taxa to meet taxonomic classification goals then are currently curated by professional staff. Thus, there is a growing need for an easy to implement tool that can generate comprehensive metabarcoding reference libraries for any bespoke locus. We address this need by reimagining CRUX from the Anacapa Toolkit and present the rCRUX package in R. The typical workflow involves searching for plausible seed amplicons (*get_seeds_local*() or *get_seeds_remote*()) by simulating *in silico* PCR to acquire seed sequences containing a user-defined primer set. Next these seeds are used to iteratively blast search seed sequences against a local NCBI formatted database using a taxonomic rank based stratified random sampling approach (*blast_seeds*()) that results in a comprehensive set of sequence matches. This database is dereplicated and cleaned (*derep_and_clean_db*()) by identifying identical reference sequences and collapsing the taxonomic path to the lowest taxonomic agreement across all matching reads. This results in a curated, comprehensive database of primer specific reference barcode sequences from NCBI. We demonstrate that rCRUX provides more comprehensive reference databases for the MiFish Universal Teleost 12S, Taberlet trnl, and fungal ITS locus than CRABS, METACURATOR, RESCRIPt, and ECOPCR reference databases. We then further demonstrate the utility of rCRUX by generating 16 reference databases for metabarcoding loci that lack dedicated reference database curation efforts. The rCRUX package provides a simple to use tool for the generation of curated, comprehensive reference databases for user-defined loci, facilitating accurate and effective taxonomic classification of metabarcoding and DNA sequence efforts broadly.

## Introduction

The fields of freshwater, estuarine and marine ecology are rapidly embracing high throughput sequencing to detect, monitor, or assess change in biological communities (Deiner et al. 2017, Takahashi et al. 2023). Fundamental to the efficacy of these molecular biomonitoring efforts, particularly metabarcoding (amplicon sequencing), is the taxonomic assignment of the sequences generated (Edgar 2018, Bik 2018). Taxonomic assignment is a complicated bioinformatic process that involves many challenges including the uncertainty around the generated sequencing data, the comparison between those data and a reference database of sequences of known origin, and the bioinformatic decisions that land on a taxonomic identification of the generated sequences (Hleap et al. 2021, Jeunen et al. 2023, Mathon et al. 2021, Edgar 2018, O’Rouke et al. 2020). Ensuring accurate taxonomic assignment is critical for the adoption of biomolecular monitoring tools including environmental DNA (eDNA) metabarcoding, microbiome, bulk metabarcoding, and gut and diet studies among others applications (Deiner et al. 2017),

Paramount to the success of taxonomic assignment is the comprehensiveness and accuracy of the reference database used to classify query DNA sequences (Banchi et al. 2020, Gold et al. 2021, Bucklin et al. 2016). Large scale barcoding of life efforts over the past three decades have provided the raw material for such reference databases (Herbert et al. 2003, Stoeckle and Herbert 2008, Costa et al. 2007, Darwin Tree of Life Project 2022) generating millions of reference barcode sequences publicly available through National Center for Biotechnology Information (NCBI), The Barcode of Life Data System (BOLD), European Nucleotide Archive (ENA), and others (Cummins et al. 2021, Ratnasingham et al. 2007, Sayers et al. 2022). Despite the incompleteness of current global reference databases across all domains of life, these large sequence repositories are constantly improving and expanding to allow for accurate identification of DNA sequences needed for a suite of ecological and public health efforts (Thompson et al. 2017, Soon et al. 2013, Manor et al. 2020, Taberlet et al. 2018, Beng and Cortlett 2020). Generating a high quality reference database from these enormous sequence repositories requires a full accounting of all orthologous sequences, the detection and removal of mislabelled sequences, and the identification of identical sequences across taxa (Curd et al. 2019, Jeunen et al. 2023, Richardson et al. 2020). Parsing and refining these large sequence repositories into curated databases that are comprehensive for specific marker sets remains a significant challenge (Jeunen et al. 2023).

Efforts to address this challenge either rely on the dedicated maintenance and curation of reference databases for specific loci of interest or are limited in efficacy because they rely on keyword searches, are too computationally demanding, or are difficult to stand up and install, relying on a suite of software dependencies. By far the most successful and widely used reference databases (e.g. Silva, PR2, UNITE, MitoFish) rely on dedicated staff and resources to maintain and update such repositories (Quast et al. 2012, Guillo et al. 2012, Kõljalg et al. 2019, Zhu et al. 2023). Given the extensive resources needed to curate and maintain such repositories, there are only a handful of such efforts representing only commonly used loci. We can not expect to have similar dedicated efforts for all metabarcoding loci. This is especially true as novel sequencing technologies allow for longer targets and more immediate *in-situ* sequencing (Zorz et al. 2023). Thus alternative reference database generating tools are needed to alleviate taxonomic assignment restrictions at the database level and fill this operational gap in the field.

A commonly utilized approach to generating reference databases relies on keyword searches (Keck et al. 2022). Such efforts are dependent on the accuracy of associated sequence metadata submitted by users. However, a lack of controlled vocabulary and metadata standards often leads to poor annotations (e.g. CO1, COI, and COX1 all describing the same cytochrome oxidase gene) which frequently limits the comprehensiveness of such reference databases (Curd et al. 2019; Jeunen et al. 2023, Hemsley et al. 2020, Porter & Hajibabaei 2018). Many tools address these specific limitations in generating comprehensive reference barcode databases for key loci like CO1 (e.g. MIDORI2, CO-ARBitrator, MARES, COInr; Leray et al. 2022, Heller et al. 2018, Arranz et al. 2020, Meglécz 2023). However, keyword search-based database generation is particularly susceptible to inadequately capturing orthologous sequences as this requires *a priori* knowledge of sequence similarities and associated metadata (e.g. MiFish 12S and microbial 16S; Gold et al. 2021). Such keyword search-based approaches are useful for a handful of widely used loci (e.g. CO1), but are not flexible enough to be applied to any metabarcoding and sequencing locus (Ahmed et al. 2019, Keck et al. 2022).

To address these limitations, a suite of reference barcode generating tools were designed based on sequence similarity instead of associated metadata to build comprehensive, curated reference databases (Jeunen et al. 2023, Richardson et al. 2020). CRUX and its counterparts, METACURATOR, Metaxa2, and CRABS, all similarly rely on a two step database generating process (Bengtsson-Palme et al. 2015,Jeunen et al. 2023, Richardson et al. 2020, Curd et al. 2019). First these tools conduct an *in silico* PCR to generate a set of “seed” sequences containing the primer regions. And since not all sequences are submitted with the primer sequences intact, these tools implement a second step which aligns these seed sequences across the large sequence repositories (e.g. GenBank, ENA, BOLD) to acquire a comprehensive set of similar sequences. Inherently, these software tools take a brute force approach to generating reference databases that acquire all orthologous sequences. Unsurprisingly, these brute force computational approaches require significant computational resources (Jeunen et al. 2023). In addition, these tools often rely on a large number of software dependencies which are difficult to install and maintain on high performance computing clusters (e.g. CRUX, Metaxa2). Together these limit the adoption and utilization of such reference database generating tools.

Here we present rCRUX, a reference database generating R package (R Core Team 2022), that relies on efficient iterative BLAST searches to sample all orthologue sequence space (Altschul et al. 1990, Ye et al. 2012), utilizing a smaller set of readily available dependencies. rCRUX provides a simple, easy to use reference database generating tool that facilitates the generation of curated, comprehensive bespoke reference databases across a diversity of users and platforms including cloud-hosted services.

## Methods

Here we build on the rationale behind the generation of locus-specific databases outlined first in Curd et al. 2019 which demonstrates that the most comprehensive databases are obtained by way of sequence similarity instead of intended taxonomic identity or sequence description (Jeunen et al. 2023, Richardson et al. 2020, Curd et al. 2019). rCRUX produces reference sequence databases in a three step process (Figure 1): 1) identification of seed sequences that match the primers of interest, 2) finding homologous and orthologous sequences to those via BLAST, and 3) dereplication of the resulting database to reduce redundancy and detect wrongly annotated sequences. This can be followed by 4) database comparison tools provided in rCRUX.

**Figure 1.**
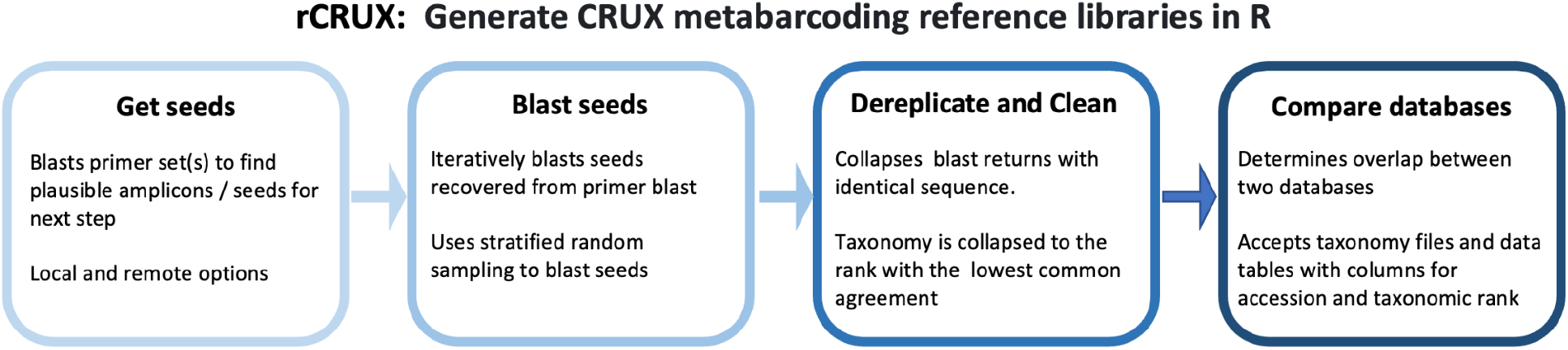
Overview of rCRUX workflow

### Installation

In order to install rCRUX onto a computer or cluster users must first download the rCRUX package and NCBI’s BLAST+ toolkit. In addition, users need a blast-formatted nucleotide database which can be downloaded directly from NCBI as well as NCBI taxonomy IDs which can be acquired using taxonomizr’s *prepareDatabase()* function (Sherrill-Mix 2019). We note that these combined databases required over 340 GB of storage as of April 2023.

### get_seeds: In silico PCR

The first step of rCRUX is an *in silico* PCR step which takes a set or sets of forward and reverse primer sequences (single or multiple forward and single or multiple reverse primers, and that can include degenerate bases) and returns possible full length barcode sequences containing forward and reverse primer matches along with taxonomic information. This step can be implemented locally through *get_seeds_local()* which uses a modified adaptation of NCBI’s primer blast or remotely through *get_seeds_remote*() which submits a web form directly to NCBI’s primer blast tool. *get_seeds_local*() avoids querying NCBI’s primer BLAST tool and thus is not subject to arbitrary throttling of remote jobs that require significant memory (Camacho et al. 2009, Ye et al. 2012).

Specifically, *get_seeds_local()* passes the forward and reverse primer sequence for a given PCR product to *run_primer_blastn()*. In the case of a non-degenerate primer set only two primers will be passed to *run_primer_blast()*. In the case of a degenerate primer set, *get_seeds_local()* will obtain all possible versions of the degenerate primer(s) (using primerTree’s *enumerate_primers*() function), randomly sample a user-defined number of forward and reverse primers, and generate a fasta file of selected primers (Cannon et al. 2016). The selected primers are then passed to *run_primer_blastn*() which queries each primer against a blast formatted database using the task *blastn_short*. This process continues until all of the selected primers are blasted. The result is an output table containing the query subject id, subject NCBI GenInfo Identifier (gi), subject accession version, number of mismatches between the subject and query, subject start base pair location, subject end base pair location, and subject NCBI taxonomic identification (taxid). These returned BLAST hits are then quality controlled to see if they generate plausible amplicons (e.g. amplify the same accession and are in the correct orientation to produce a PCR product). These hits are further filtered for user-specified length and number of mismatches. Lastly, taxonomy is appended to these filtered hits using *get_taxonomy_from_accession()* (Sherrill-Mix 2019, Zhang et al. 2017, Zeileis et al. 2005).

Alternatively, *get_seeds_remote()* passes the forward and reverse primer sequence along with user-specified taxid(s) of target organism(s) and databases through *iterative_primer_search* to NCBI’s primer blast tool (Ye et al. 2012). Degenerate primers are converted into all possible non-degenerate sets and a user-defined maximum number of primer combinations is passed to the API. Multiple taxids are searched independently, as are multiple databases (e.g. c(‘nt’, ‘refseq_representative_genomes’). A primer search is then conducted on each resultant combination using *modifiedPrimerTree_Functions* which is a modified version of primerTree’s *primer_search*() and primerTree’s *parse_primer* to query NCBI’s primer BLAST tool, filter the results, and aggregate them into a single data.frame (Cannon et al. 2016). These hits are further filtered for user-specified length and number of mismatches. Lastly, taxonomy is appended to these filtered hits.

The result of the two *get_seeds_\** functions is a fasta and taxonomy file of sequences containing the specified primer sequences along with summary statistics of accessions and hits returned.

### blast_seeds: Building Comprehensive Databases of Similar Sequences

*blast_seeds()* takes the output from *get_seeds_local()* or *get_seeds_remote()* and iteratively blasts seed sequences using stratified random sampling of a given taxonomic rank making sure that at least one representative of each rank enters the iterative blast process (default rank is genus). The randomly sampled subset of seeds is then formatted into a multi-line fasta and run through *blastn* to recover all similar sequences based on user defined sequence similarity parameters (e.g., percent identity, evalue, query length; Camacho et al. 2009). The resulting blast hits are then de-replicated by accession with only the longest read per accession retained in the output table. All of the subset seeds and seeds recovered through the blast process are then removed from the seeds dataframe, reducing the number of sequences to be blasted. This stratified random sampling process is repeated until there are fewer seed sequences remaining than the max_to_blast parameter, at which point all remaining seeds are blasted. The final aggregated results are cleaned for multiple blast taxids, hyphens, and wildcards and are then appended with taxonomy.

Importantly, we note that the blast databases downloaded from NCBI’s FTP site utilizes representative accessions where identical sequences have been collapsed across multiple accessions even if they have different taxids. Here we identify representative accessions with multiple taxids (Katz et al. 2021), and unpack all of the collapsed accessions to allow for the identification of lowest common agreed taxonomy for each representative accession. We do not identify or unpack representative accessions that report a single taxid.

### derep_and_clean_db: Quality Control and Curation of Reference Database

The final step of rCRUX, *derep_and_clean_db()*, takes the output from *blast_seeds()* and conducts quality control and curation de-replicates the dataset to identify representative sequences. First, all sequences with NA taxonomy for phylum, class, order, family, and genus are removed from the dataset because they typically represent environmental samples with low value for taxonomic classification and are stored separately. Next, all sequences with the same length and composition are collapsed to a single database entry, where the accessions and taxids (if there are more than one) are concatenated. The sequences with a clean taxonomic path (e.g. no ranks with multiple entries) are saved. In contrast, sequences with multiple entries for a given taxonomic rank are processed further by removing NAs from rank instances with more than one entry (e.g. “Chordata, NA” will mutate to “Chordata”). Any remaining instances of taxonomic ranks with more than one taxid are reduced to NA (e.g. species rank “*Badis assamensis, Badis badis*” will mutate to “NA”, but genus rank will remain “*Badis*”). Finally, the resulting taxonomic paths are synonymized to the lowest taxonomic agreement. Lastly, the above cleaned and dereplicated sequences are used to generate a fasta file and taxonomy file of representative NCBI accessions for each sequence.

### compare_ref_db

*Exploring Overlap and Mismatches Between Two Reference Databases* We provide an additional function to compare the overlap and mismatches between any two reference databases. This function provides a summary table, generates a venn diagram of overlapping accessions and species, and creates a krona plot of unique taxa to each reference database (Gao et al. 2021; McMurdie and Holmes 2013, Pauvert 2020).

### Shiny Application

We further provide an R Shiny application to facilitate rCRUX accessibility. *Shiny* apps are click-based graphical user interface (GUI) programs that allow users not familiarized with R statistical software to interact with R products. The workflow outlined above requires the user to provide a primer pair of interest to generate a database of orthologous sequences and their associated taxonomies. Between input and output, users have to download two databases from the *taxonomizr* package that links NCBI’s accessions with taxids (Sherrill-Mix 2019); and taxids with higher and lower taxonomic ranks to establish the full lineage of a taxa. Furthermore, an improved performance is achieved if all BLAST operations are conducted locally and not limited by NCBI’s servers (Camacho et al. 2009). As of April 2023, over 340 GB of information have to be downloaded, which limits the real-world utility of any software application. With this in mind, we have created an interactive *Shiny* app that will allow all large file downloads to occur remotely by a host computer, be that a dedicated server in a laboratory, a computational cluster at a research institution, or an open-access portal built on cloud computing. Thus, the preparation and computational burdens of the processes are greatly reduced for the end user of the databases, who only need to provide the DNA sequences of the desired primers.

### Benchmarking

We first benchmark the efficacy of the seed generating step and sequence accumulating step of several database building tools (as published in Jeunen et al. 2023) using the following barcode markers: MiFish Universal Teleost (MiFish; Miya et al. 2015), Taberlet trnl (trnl; Taberlet et al. 1991, Taberlet et al. 2007), and fungal ITS (FITS;Ihrmark et al. 2012, White et al. 1990). We built rCRUX databases using the optimized parameters outlined in the Supplemental Methods (Figures S1-S8). The trnl and FITS databases were made using the NCBI nt blast database (see Supplemental Methods for details), and we made two rCRUX MiFish databases; one using the NCBI nt blast database, and one using NCBI nt blast database and an additional custom blast database comprised of all Actinopterygii mitogenomes (see Supplemental Methods for details). The expanded database is used for all comparisons below. rCRUX reference database statistics are presented in Table 1 and Supplemental Table 3).

We benchmarked the get_seeds_local() *in silico PCR function* with the ECOPCR (Ficetola et al. 2010) databases created by Jeunen et al. 2023. We next benchmarked the databases generated by the *blast_seeds*() by comparing the overlapping ncbi accessions and taxonomic content of the rCRUX databases against the CRABS (All Markers), METACURATOR (All Markers), ECOPCR (All Markers), RESCRIPt (MiFish, trnl) presented in Jeunen et al. 2023 (Richardson et al. 2020, Robeson et al. 2021) and a CRUX generated reference databases (Gold et al. 2021, MiFish only). We note that rCRUX was generated with NCBI nt database downloaded December 2022 (see Supplemental Methods) and thus contains more sequences than the compared reference databases presented in Jeunen et al. 2023 and Curd et al. 2019.

We then generated rCRUX reference databases for 16 metabarcoding primer sets (Table 1) and made version-controlled, DOI accessions available on the GitHub to provide comprehensive curated reference databases for a suite of bespoke metabarcoding loci that lack dedicated reference databases.

## Results

The *get_seeds_local*() *in silico PCR* consistently captured a greater number of species than ECOPCR across the MiFish 12S Universal Teleost, trnl, and FITS loci (Figure 1a,b,c). *get_seeds_local()* captured 95.2%, 86.7%, and 63.6% of species that ECOPCR captured while also capturing an additional 6,715 (40% of total species), 16,242 (74%), and 39,792 (56%) species respectively. Overlap in species captured ranged from 57% for the MiFish locus to 23% for the trnl locus (FITS was 28%).

We further demonstrate that rCRUX reference databases consistently capture a greater number of accessions and species for MiFish (Figure 2), trnl (Figure 3), and FITS (Figure 4) loci across comparable benchmark reference databases.

**Figure 2.**
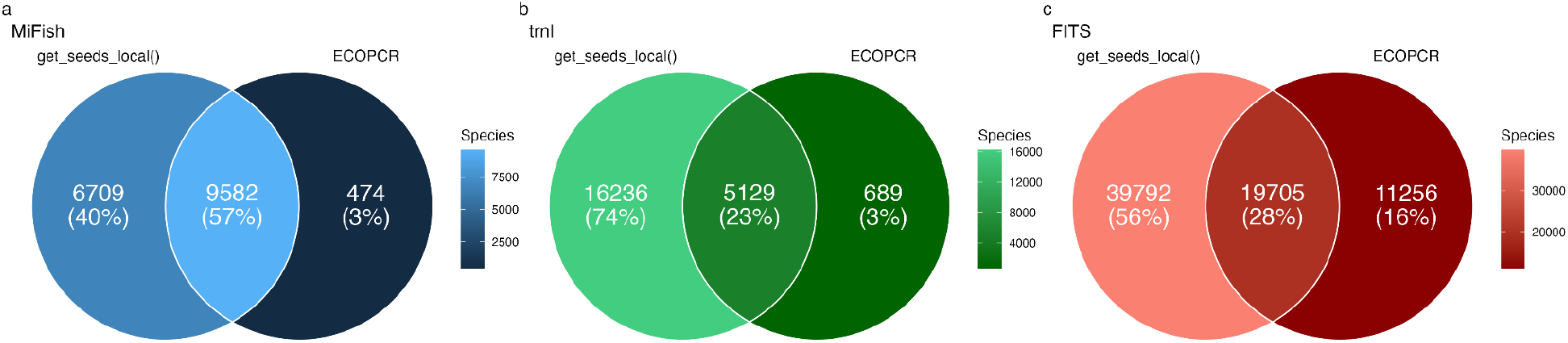
rCRUX *get_seeds_local*() *in silico* PCR comparison with ECOPCR. Comparison of number of species captured by rCRUX *get_seeds_local()* and OBITOOLS ECOPCR *in silico* PCR tools for a) MiFish 12S Universal Teleost, b) trnl, and c) fungal ITS (FITS) loci.

**Figure 3.**
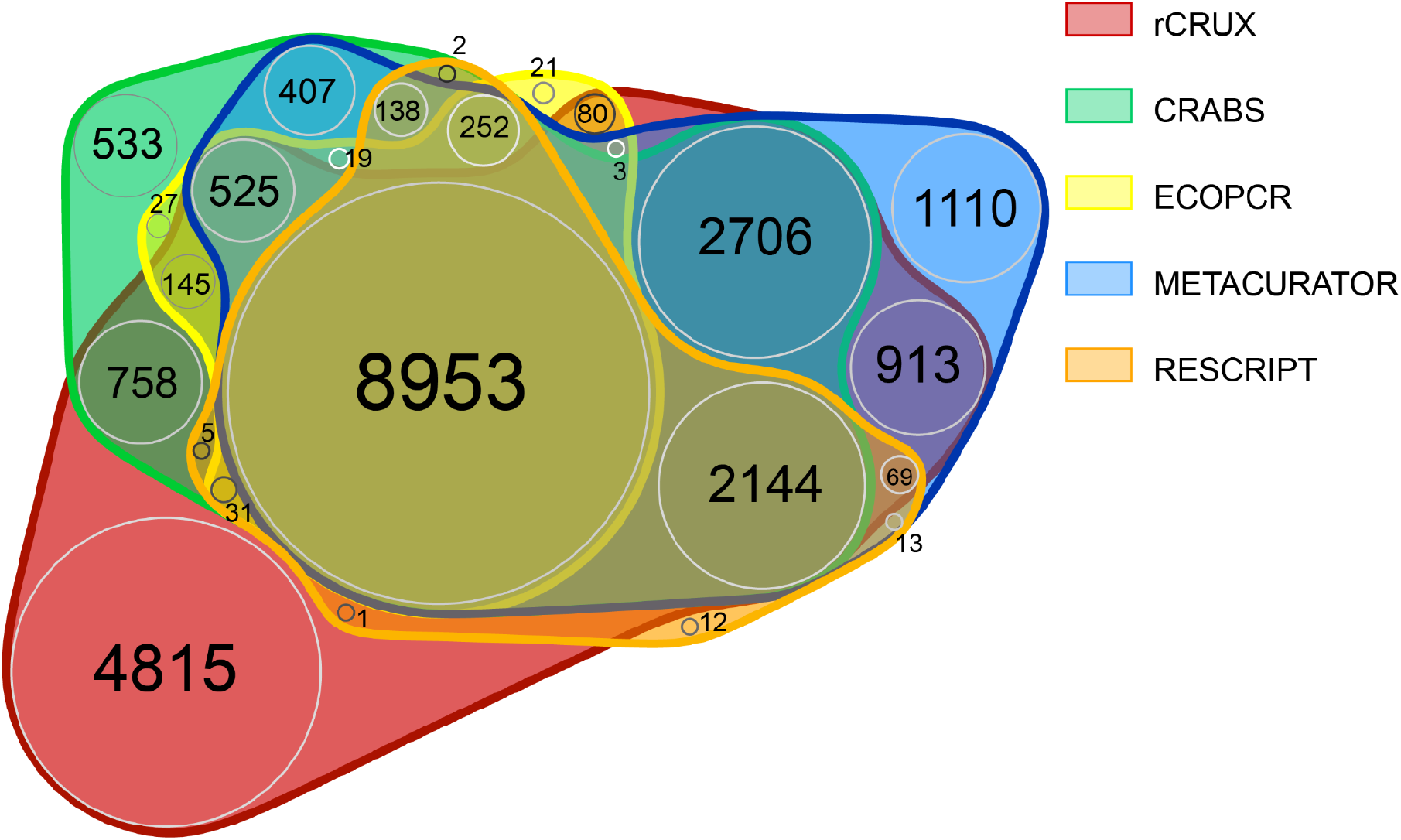
MiFISH rCRUX *blast_seeds*() database comparison with CRUX, CRABS, ECOPCR, METACURATOR, and RESCRIPt databases.

**Figure 4.**
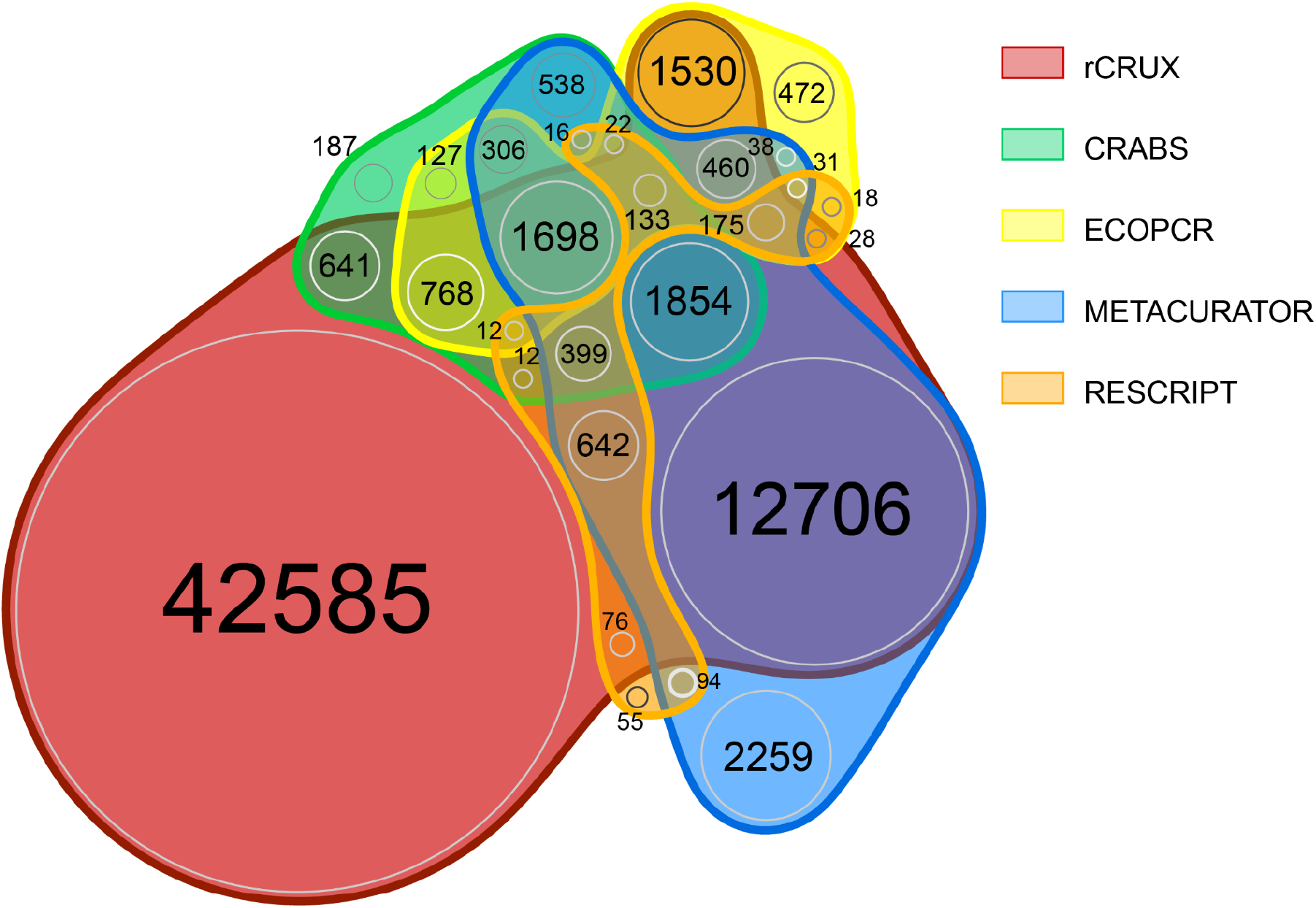
trnl rCRUX *blast_seeds*() database comparison with CRABS, ECOPCR, METACURATOR, and RESCRIPt databases.

For the MiFish reference database, only 32.5% of all species (n=7,933) shared across the 6 reference databases. Each reference database had unique sequences that were not shared with any other database (range: 12 - 4,815). rCRUX captured 86.6% (n=21,148) of all species observed across the MiFish reference databases. rCRUX uniquely had 14.2% (n=3,459) of all species observed and 6% (n=1,356) shared only with the CRUX reference database.

For the trnl reference database, only 133 species out of 67,882 species shared across the 5 trnl reference databases. Each reference database had unique sequences that were not shared with any other database (range: 55 - 42,585). rCRUX captured 93.9% (n=63,719) of all species observed across the trnl reference databases. rCRUX uniquely had 62.7% (n=42,585) species not observed in any other database and 18.7% (n=12,706) shared only with the METACURATOR database (Figure 3).

For the FITS reference database, only 4.6% of all species (n=11,121) were shared across the 4 reference databases. Each reference database had unique sequences that were not shared with any other database (range: 920 - 177,272). rCRUX captured 94.6% (n=228,825) of all species observed across the FITS reference databases. rCRUX uniquely had 73.3% (n=177,272) species (Figure 3).

Limiting the seeds and database generation output comparisons to only Eukaryotic reads had minimal effect on the results (Supplemental Figures S9-12). We also note that the rCRUX databases were generated after the other databases, however they include the majority of species captured by compared methods. Together, these results benchmark rCRUX favorably against CRABS, METACURATOR, ECOPCR, RESCRIPt, and CRUX across a diversity of metabarcoding loci.

### rCRUX databases

We successfully generated a total of 16 reference databases (Table 2) for a suite of bespoke metabarcoding primer sets. Sizes of these reference databases ranged from 576 to 1,371,297 accessions and 206 (160 Unique reads) to 228,874 (138,089 unique reads) species.

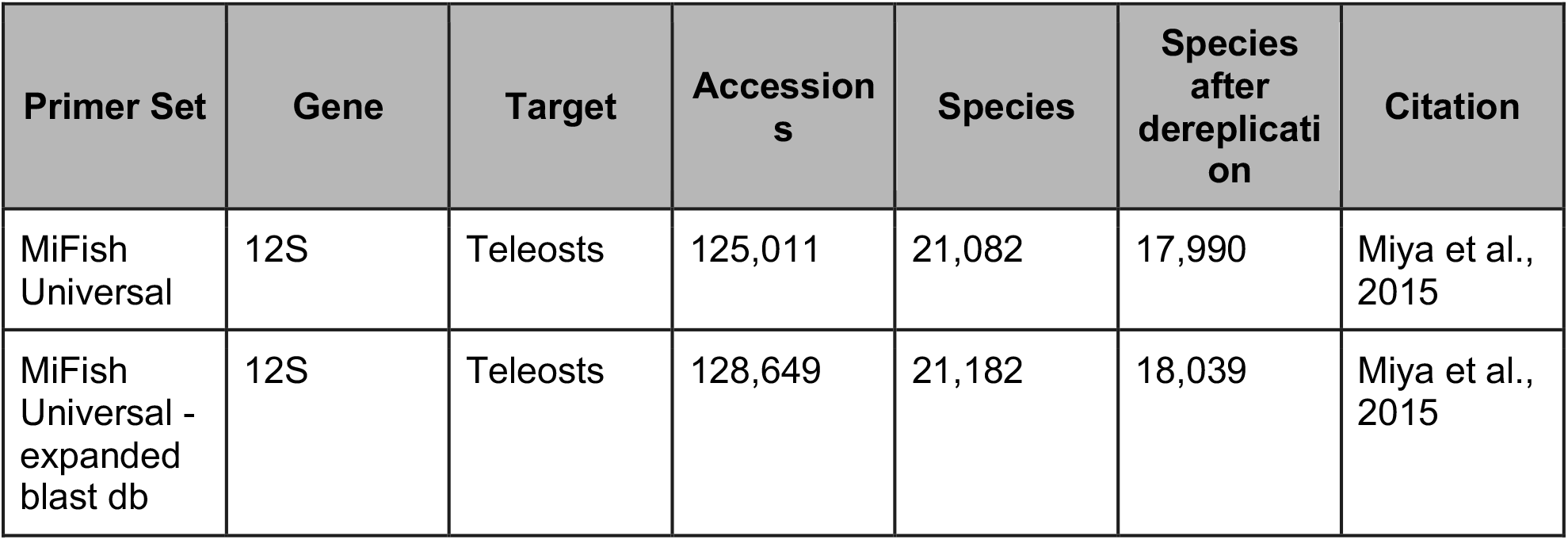
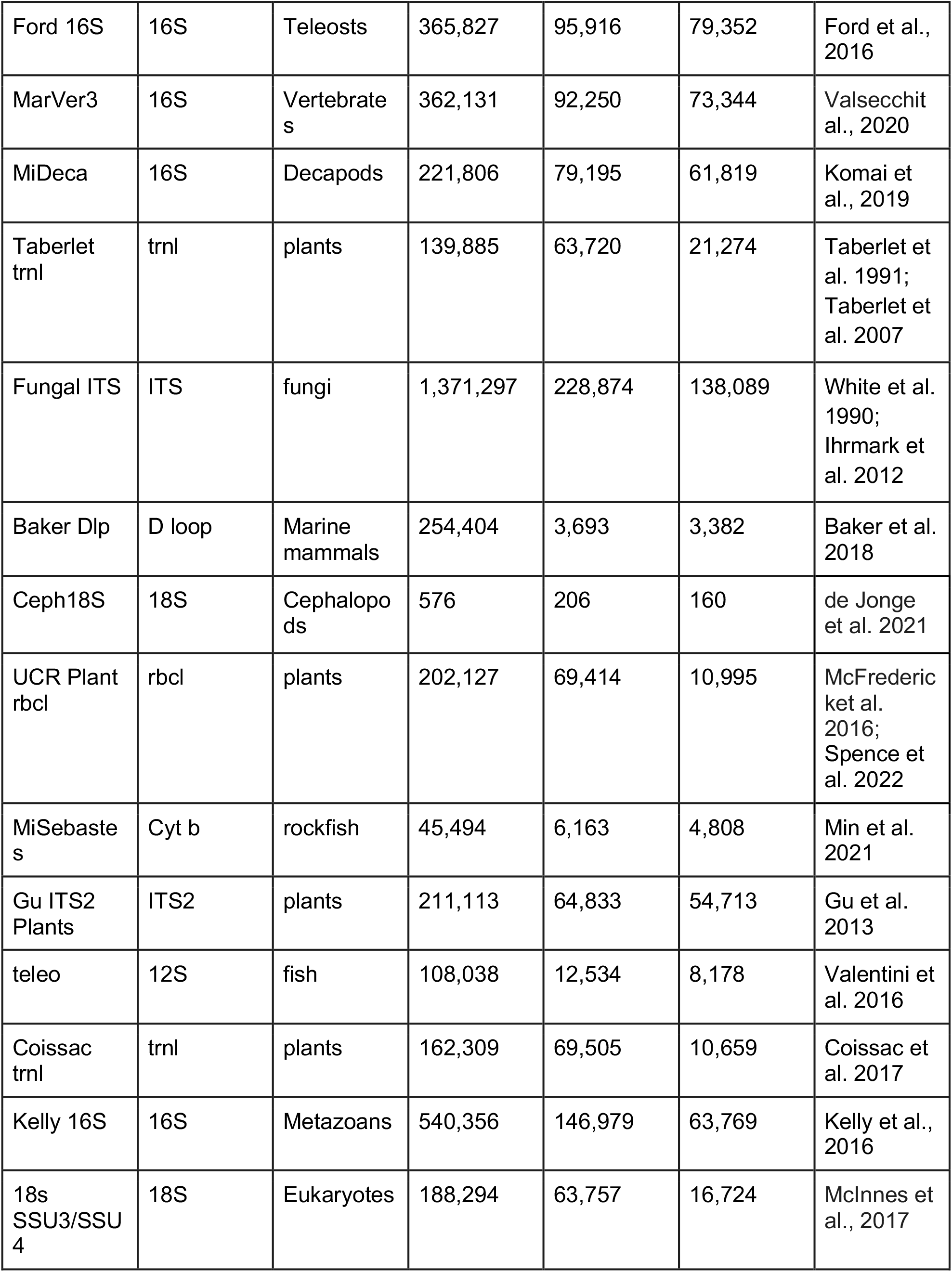

| Primer Set | Gene | Target | Accessions | Species | Species after dereplication | Citation |
| --- | --- | --- | --- | --- | --- | --- |
| MiFish Universal | 12S | Teleosts | 125,011 | 21,082 | 17,990 | Miya et al., 2015 |
| MiFish Universal - expanded blast db | 12S | Teleosts | 128,649 | 21,182 | 18,039 | Miya et al., 2015 |
| Ford 16S | 16S | Teleosts | 365,827 | 95,916 | 79,352 | Ford et al., 2016 |
| MarVer3 | 16S | Vertebrates | 362,131 | 92,250 | 73,344 | Valsecchi et al., 2020 |
| MiDeca | 16S | Decapods | 221,806 | 79,195 | 61,819 | Komai et al., 2019 |
| Taberlet trnl | trnl | plants | 139,885 | 63,720 | 21,274 | Taberlet et al. 1991; Taberlet et al. 2007 |
| Fungal ITS | ITS | fungi | 1,371,297 | 228,874 | 138,089 | White et al. 1990; Ihrmark et al. 2012 |
| Baker Dlp | D loop | Marine mammals | 254,404 | 3,693 | 3,382 | Baker et al. 2018 |
| Ceph18S | 18S | Cephalopods | 576 | 206 | 160 | de Jonge et al. 2021 |
| UCR Plant rbcl | rbcl | plants | 202,127 | 69,414 | 10,995 | McFrederick et al. 2016; Spence et al. 2022 |
| MiSebastes | Cyt b | rockfish | 45,494 | 6,163 | 4,808 | Min et al. 2021 |
| Gu ITS2 Plants | ITS2 | plants | 211,113 | 64,833 | 54,713 | Gu et al. 2013 |
| teleo | 12S | fish | 108,038 | 12,534 | 8,178 | Valentini et al. 2016 |
| Coissac trnl | trnl | plants | 162,309 | 69,505 | 10,659 | Coissac et al. 2017 |
| Kelly 16S | 16S | Metazoans | 540,356 | 146,979 | 63,769 | Kelly et al., 2016 |
| 18s SSU3/SSU4 | 18S | Eukaryotes | 188,294 | 63,757 | 16,724 | McInnes et al., 2017 |

## Discussion

We successfully demonstrate that rCRUX generates comprehensive, curated reference databases for user-defined metabarcoding loci of interest. We benchmarked rCRUX against similar reference database generating tools, consistently capturing the majority of accessions and species present in those databases as well as thousands of additional species and accessions not found in CRABS, METACURATOR, ECOPCR, RESCRIPt, and the original implementation of CRUX (Ficetola et al. 2010, Jeunen et al. 2023, Curd et al. 2019, Richardson et al. 2020, Robeson et al. 2021). We generated 16 reference databases for bespoke metabarcoding loci of interest, providing important bioinformatic resources for the broader metabarcoding community. The rCRUX R package presented here provides a valuable tool for the generation and curation of reference databases, enhancing the accuracy, utility, and reproducibility of taxonomic assignment of DNA sequences broadly.

### Benchmarking rCRUX reference databases

rCRUX generated reference databases were consistently more comprehensive than other leading databases (Figures 2-5, Supplementary Figures S9-12). This is partially driven by the efficacy of *get_seeds_local*(), which captured more species and accessions than ECOPCR, serving as a more efficient *in silico* PCR simulator (Curd et al. 2019; Ficetola et al. 2010) (Figure 1). At the core of *get_seeds_local*() is the NCBI primer blast tool which is a widely used, well benchmarked, and reproducible tool for the testing of primer sets (Ye et al. 2012, Cannon et al. 2016, Hleap et al. 2021). In addition, we demonstrate that our iterative blasting approach did not impair the efficacy of the *blast_seeds()* step as originally implemented in CRUX, resulting in faster run times and comparably comprehensive reference databases (Figures 2-5; supplemental results). Previous research has highlighted the value of more comprehensive reference databases for improved taxonomic assignment (Keck et al. 2022, Gold et al. 2021, Jeunen et al. 2023, Curd et al. 2019, Richardson et al. 2020). Thus, the greater diversity and breadth of species and accessions captured in rCRUX generated reference databases provides an important tool for improving taxonomic classification.

**Figure 5.**
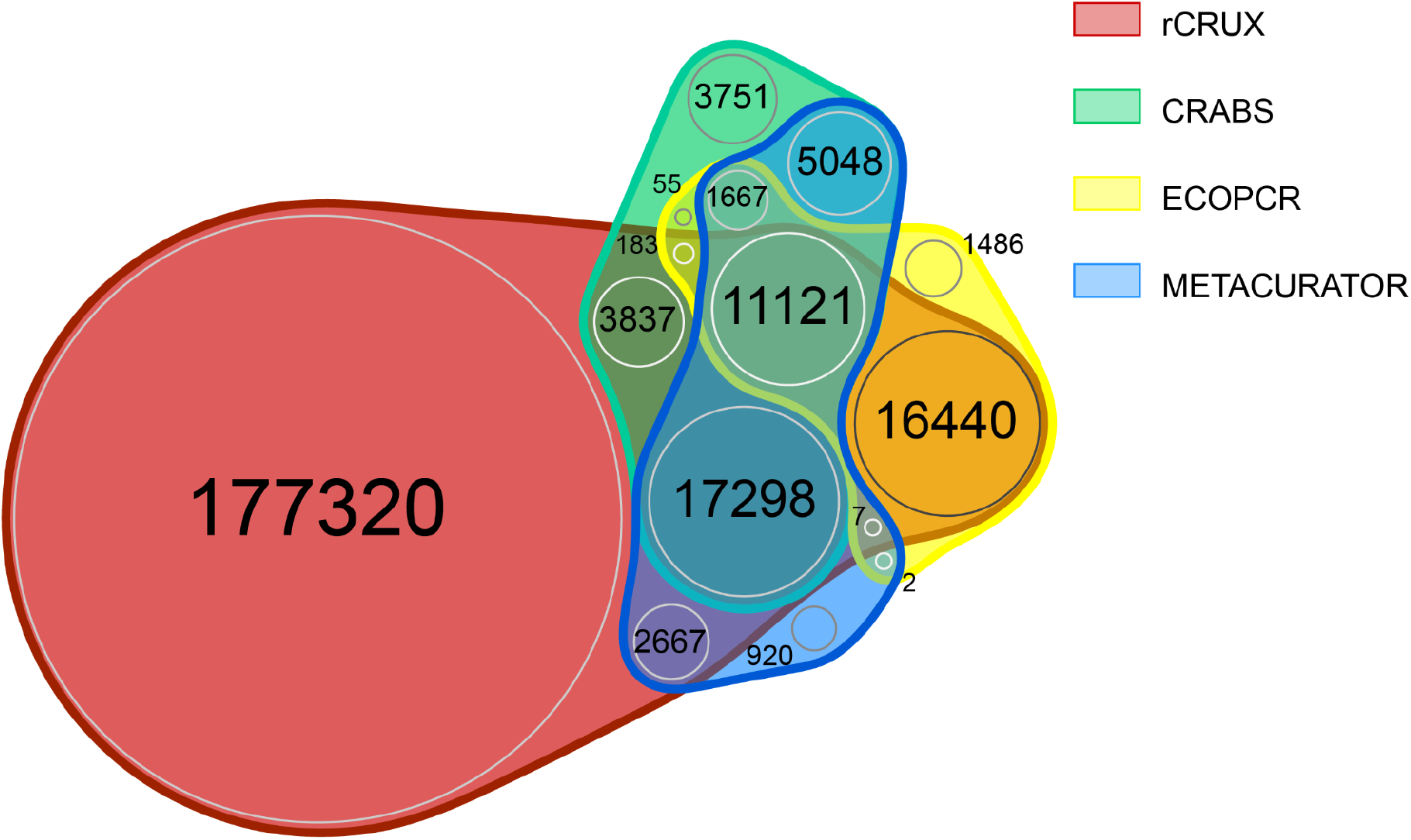
FITS rCRUX *blast_seeds*() database comparison with CRUX, CRABS, ECOPCR, METACURATOR, and RESCRIPt databasess.

Interestingly, we found that no reference database was completely comprehensive as each generated reference database tested had unique accessions and species (Figures 2-5). These patterns are consistent with previous reference database comparisons and highlight the inherent difficulty of capturing all relevant accessions for a given metabarcoding locus of interest (Jeunen et al. 2023, Curd et al. 2019, Richardson et al. 2020). Although each of the tools compared here share an underlying strategy for creating comprehensive reference databases for loci of interest, their implementation is different, resulting in distinct subsets of captured reference sequences from public DNA repositories.These results highlight the challenge of making reproducible reference databases. However, despite these differences across reference database generators, rCRUX captured the vast majority of all species and accessions captured across the tested reference databases, failing to capture at most ∼5% of sequences captured by another database. Future reference database generating efforts may seek to employ multiple distinct generating strategies and combine results to obtain the most comprehensive reference database possible.

Importantly, we demonstrate that rCRUX is nearly perfectly reproducible across subsequent runs of MiFish Teleost rCRUX databases (n=10 runs, coefficient in variation of returned accessions <0.01%, see supplemental results and Supplementary Table 3), providing high confidence in rCRUX reference databases.

### FAIR Reference Database Generation

Access to reliable, reproducible, comprehensive, and curated reference databases is critical for improving taxonomic assignment of DNA sequences (Keck et al. 2022). However, to date, curated, metabarcode specific reference database generating tools have not fully adhered to the Findable, Accessible, Interoperable, and Reproducible data management principles (Wilkinson et al. 2016). The generation and curation of reference barcode databases is time and labor intensive and requires substantial computational resources and bioinformatic expertise which often limits interoperability and reproducibility across users. Together, this severely limits our ability to generate reference databases quickly and efficiently and limits the number of researchers and scientists who can build the repositories needed to assign taxonomy to DNA sequences (Curd et al. 2019, Shea et al. 2023).

However, recent advances in reference database generating tools open the door to a broader community of practice in generating reference databases (Jeunen et al. 2023). Given the ubiquity of R users in the molecular biology and ecology fields, rCRUX provides a powerful tool that is straightforward and relatively easy to implement on any computing environment. By providing researchers with an accessible reference database generating tool, we hope to alleviate the difficulties of building and updating reference databases. Thus the ability to generate user-specific barcode reference databases will enhance metabarcoding, eDNA, microbiome, and DNA classification research efforts broadly.

One of the motivations for making simple and easy to install, update, and maintain reference database generating tools was to increase access to these resources across the molecular biology and ecology fields. However, limitations in the utility of reference database generating software still remain, particularly the scale of computational resources needed.

Although the iterative blast implementation of rCRUX reduces computational needs compared to the previous iterations of CRUX, the rCRUX databases presented here still relied on high performance computing (each run was given a maximum allotment of 250GB of RAM, 40 cores, and one week of run time on the University of Vermont - Vermont Advanced Computing Cluster). Even with such computational power, efforts to generate larger reference databases with greater number of available reference barcodes (e.g. CO1, microbial 16S and 18S) were unsuccessful because of a lack of available computational resources to meet the scale of available sequences. Fortunately such markers have dedicated efforts to curate comprehensive and accurate reference databases (e.g. MIDORI2, MARES, Silva, PR2) and researchers can easily access such databases (Leray et al. 2022, Arranz et al. 2020, Quast et al. 2012, Guillo et al. 2012). Researchers often lack access to computational resources, particularly in developing nations where biodiversity is often the highest and the need for DNA-based taxonomic classification is greatest (Barber et al. 2014, Asase et al. 2022., Johnson et al. 2022). As cloud computing and high performance computing resources continue to become increasingly cost effective, we hope rCRUX and similar reference database generating tools will become more accessible (Thompson & Thielen 2023). We note that rCRUX can be successfully implemented on a personal laptop with a 1 TB hard drive, 16 GB of RAM, and 8 cores, given parameters and markers that require fewer computational resources. Importantly, we designed rCRUX to be highly scalable and easy to install through R in any compute environment, allowing for adoption in future cloud computing efforts in which rCRUX could be served to a wide audience like NCBI primerTools or BLAST.

However, to specifically help address issues of access to comprehensive reference databases in the short term, we provided 16 reference databases for commonly used or emerging metabarcoding loci. These databases will be updated and curated at least annually with a unique DOI, providing important genetic resources to the broader DNA sequencing community including those that lack access to such computational infrastructure. Future efforts will be made to grow the list of available databases as future loci become available and widely adopted.

Lastly, we demonstrate the reproducibility of rCRUX, allowing for users to make identical databases from the same starting parameters and sequence repositories (Supplemental Table 2). Providing a reproducible and stable tool for the generation of barcode reference databases ensures high quality genetic resources that adhere to FAIR principles.

### Broader Applications of rCRUX

The most immediate application of rCRUX is the generation of reference databases to support taxonomic assignment of metabarcoding from high throughput sequencing. However, the utility of rCRUX allows for reference databases to be generated on any blast formatted database, directly supporting improved taxonomic assignment of a broad range of DNA sequence applications. For example, this allows for the curation of reference barcodes from full or partial length mitogenomes (See supplemental methods), supporting long read sequencing taxonomic assignment applications (Johri et al. 2019, Ramon-Laca et al. 2022). In addition, rCRUX can be used for building nuclear DNA-based reference databases from whole genome and transcriptome sequences. Such efforts could be used to develop population genetic and eRNA specific reference databases for a diversity of biomonitoring applications (Simon et al. 2019, Greco et al. 2022, Adams et al. 2019, Sigsgaard et al. 2020, McKinney et al. 2022).

Importantly, reference database generating tools like rCRUX provide a valuable resource for designing, validating, and comparing potential metabarcoding targets for a specific research question (Hleap et al. 2021, Mathon et al. 2021, Edgar 2018). For example, if researchers are deciding which fish metabarcoding loci to use for a given project and have a known target species list (Jerde et al. 2021), rCRUX can be used to conduct an *in silico* comparison of primer set efficacy. This can be accomplished by first generating rCRUX reference databases for each potential locus and then by cross referencing the taxonomic resolution of each database against the target taxa list (Gold et al. 2021). Furthermore, the comparison and curation tools provided in the R package allows for the direct comparison of multiple reference databases, serving as a resource for evaluating the relative performance of reference databases on taxonomic assignment. Previous research has demonstrated the value of these kinds of *in silico* validation and benchmarking approaches for improved taxonomic classification of DNA sequences (Edgar 2018, Curd et al. 2019, Gold et al. 2021, Jeunen et al. 2023). Thus rCRUX provides a simple, cost effective tool for informing scientists and resource managers on the efficacy of taxonomic assignment during the design and development of biomolecular monitoring efforts.

### Complimentary Packages to rCRUX

The rCRUX package provides important novel utility to the wide suite of reference database managing packages available. We note that such packages can be used in concert to achieve improved reference database management and efficacy. For example, the refdb R package provides a suite of complementary tools that can be used to merge BOLD and GenBank databases which could provide improved blast-formatted nucleotide databases (Keck et al. 2023). In addition, refdb provides a suite of tools to visualize and summarize output reference databases (Keck et al. 2023). Similar utilities to merge GenBank, EMBL, and BOLD databases are available through CRABS, MARES, and RESCRIPt and can be used to generate a more comprehensive starting blastDB database, particularly for CO1 genes (Arranz et al. 2020, Jeunen et al. 2023, Robeson et al. 2021). In addition, CRABS and MARES also provide tools to output datasets in a greater diversity of formats for use in additional taxonomic classifiers beyond Anacapa and Qiime2 (Curd et al. 2019, Boylen et al. 2019). The comprehensiveness of rCRUX databases can also be leveraged and used as input into GAPeDNA to better conduct gap analysis for a given locus and target taxa in a specific study region (Marques et al. 2021). Similarly, researchers submitting ‘Omics data to Ocean Biogeographic Information System (OBIS) and complying with Darwin Core standards can use the World Register of Marine Species taxonomy and can readily convert rCRUX taxonomy using the worrms R package (Meyer et al. 2021, Berry et al. 2021, Costello et al. 2013, Chamberlain 2020, Grassle and Stocks 1999). Thus rCRUX provides an important complementary tool to the suite of available reference database management software.

## Conclusion

Ultimately, rCRUX provides a powerful, reproducible, and reliable tool for the generation of comprehensive and curated reference databases for any genetic loci of interest. By providing users with a simple and accessible reference database generating R package, rCRUX will ease taxonomic classification as well as validation and benchmarking for bespoke and novel primer sets. Improved ease of implementation over previous iterations of CRUX as well as a suite of 16 publicly available version-controlled reference databases provide important genetic resources without the need of significant computational resources, facilitating access and adoption of high quality reference databases and database generating tools to a broad range of users.

## Supporting information

Supplemental Methods and Results

## Acknowledgements

This work benefited from the amazing input of many including Lenore Pipes, Sarah Stinson, Gaurav Kandlikar, Maura Palacios Mejia, Ryan Kelly, and Kim Parsons. We want to especially acknowledge the late, great Jesse Gomer, coding extraordinaire, rCRUX co-conspirator, and dear friend who tragically passed away before rCRUX was completed. None of this would be possible without Jesse’s endless inspiration, creativity, ingenuity, and generosity.

## Data Availability

The rCRUX package and 16 generated reference databases are available at https://github.com/CalCOFI/rCRUX. Data and code for analysis and figures will be uploaded upon acceptance of the manuscript.

