## Supplemental Methods and Results for "rCRUX: A Rapid and Versatile Tool for Generating Metabarcoding Reference libraries in R"

**Supplemental Materials**

### **Supplemental Methods**

*Databases*

All databases presented were built using an NCBI nt blast database (ftp://[ftp.ncbi.nlm.nih.gov/blast/db/nt](http://ftp.ncbi.nlm.nih.gov/blast/db/nt)) downloaded on December 22, 2022. The expanded 12S database was built by downloading all Actinopterygii mitochondrial genomes (available April 20, 2023) and building a blast database using the blast+ 2.13.0 [makeblastdb](https://rdrr.io/github/mhahsler/rBLAST/man/makeblastdb.html) function (ftp://[ftp.ncbi.nlm.nih.gov](http://ftp.ncbi.nlm.nih.gov/blast/db/nt)/blast/executables/blast+). The taxonimizr database was built with the ncbi taxdump (<https://ftp.ncbi.nlm.nih.gov/pub/taxonomy/>) available on December 12, 2022.

*Parameter tests*

We preformed optimization tests of get_seeds_local() and blast_seeds() parameters using two distinct metabarcode markers were used to optimize rCRUX run parameters: MiFish Universal 12S (Miya et al., 2015) and rbcL (McFrederick and Rehan, 2016). Both markers target at least a phylum of organisms (Chordata and Streptophyta) with tens of thousands of available reference sequences and are thus representative of many common metabarcoding loci. For get_seeds_local() and blast_seeds(), we tested the effects of combinations of the following parameters on the number of sequences and unique taxonomic returned by each function: max number of blast alignments (“align”; '100000000', '10000000', and '1000000'), evalue ('3e+06', '3e+07', '3e+08'), percent identity of primer query to subject (“perID”; 70, 90, 100), percent coverage of the primer query to subject (“coverage”; 70, 90, 100), and rank (“species”, “genus”, “family") for blast_seeds() only. All tests were performed on the Vermont Advanced Computation Core and allotted a maximum of 256 GB of memory and 40 Cores. For get_seeds_local() we compared the difference between filtered and unfiltered sequence returns and the number of taxonomic ranks for filtered sequence returns. For blast seeds we compared the total number of sequence returns and the number of taxonomic ranks in R.

*Database Comparisons Filtered for Eukaryotes*

We repeated the comparisons presented in the Results section filtering out all non-eukaryotic taxa.

*Reproducibility*

The function blast_seeds() uses a random stratified sampling approach to minimize the computation resources required to blast all seeds in from output of the get_seeds functions. To estimate the effect of this sampling approach we made 10 replicate blast_seeds() databases for the MiFish Universal 12S library using the same get_seeds_local() output. Note that get_seeds_local() will always produce the same output given the same parameters and query databases. Also note that balst_seeds() will produce the same database given the same parameters and query databases and if the parameter random_seed is set to an identical value between runs. The commands used to build the databases are as follows:

forward_primer_seq = "GTCGGTAAAACTCGTGCCAGC", reverse_primer_seq = "CATAGTGGGGTATCTAATCCCAGTTTG"

*get_seeds_local*(forward_primer_seq, reverse_primer_seq, metabarcode_name, output_directory_path, accession_taxa_sql_path, blast_db_path, align = '1000000', perID = 70, coverage = 70, evalue = '3e+06', num_threads = 40, max_to_blast = 1, minimum_length = 170, maximum_length = 250)

*blast_seeds*(seeds_output_path, Blast_db_path, accession_taxa_sql_path, output_directory_path, Metabarcode_name, expand_vectors = TRUE, minimum_length = 140, maximum_length = 250, num_threads = 40, max_to_blast=100, align='1000000', coverage=70, perID=70, evalue = '3.00E+07', rank = 'genus')

*Database generation*

We made 16 rCRUX databases representing metabarcodes that target a range of taxonomic groups (single phyla to several domains (e.g. NCBI rank “superkingdoms”). All were run with the following parameters for get_seeds_local(): align = '10000000', perID = 70, coverage = 70, evalue = '3e+06', num_threads = 40, max_to_blast = 1. All were run with the following parameters for blast_seeds(): num_threads = 40, max_to_blast=100, align='1000000', coverage=70, perID=70, evalue = '3.00E+07', rank = 'genus', and random_seed =23. The minimum and maximum length, barcode sequence, and blast_seeds() max to blast parameter values are given in SupplementaryTable 1.

For large returns databases (e.g. Machida 18S, 18S SSU3/5), we used R (R Core Team 2022) library dplyr (Wickham et al., 2023) to filter the get_seeds_local() output into several smaller datasets (e.g. Prokaryotes, samples with NA, Metazoans, Fungi, and Viridiplantae) and ran blast_seeds() on those databases. We combined Summary.csv tables, fasta and taxonomy files, and removed duplicate returns. These results are reported in Table 1 and Supplementary Table 3.

### **Supplemental Results**

*Parameter Optimization*

We found a significant effect of parameter choice on the number of observed returned accessions and species for *get_seeds_local*() and *blast_seeds*() across both MiFish and trnl markers.

*MiFish marker*

We found a significant effect of percent coverage and percent identity on the number of returned accessions and species observed from *get_seeds_local*() on the MiFish locus (ANOVA, P <0.0001, Figures S1 &S2, Table S4). Percent cover and percent identity had a significant effect on the number of returned accessions and species (ANOVA, P <0.0001). The max number of blast alignments and e-value had no significant effect on the number of returned accessions or species.

We observed the highest return accessions and species using percent coverage of 70, percent identity of 70, e-value 3e+7, and max number of blast alignments = '100,000,000'. We note that changing the max number of blast alignments = '10,000,000' produced nearly as many accessions and species as the above parameters, but was much less computationally intensive. Tests using parameters of e-value of 3e+8 timed out from a lack of computational resources more frequently than any other method (Table S4).

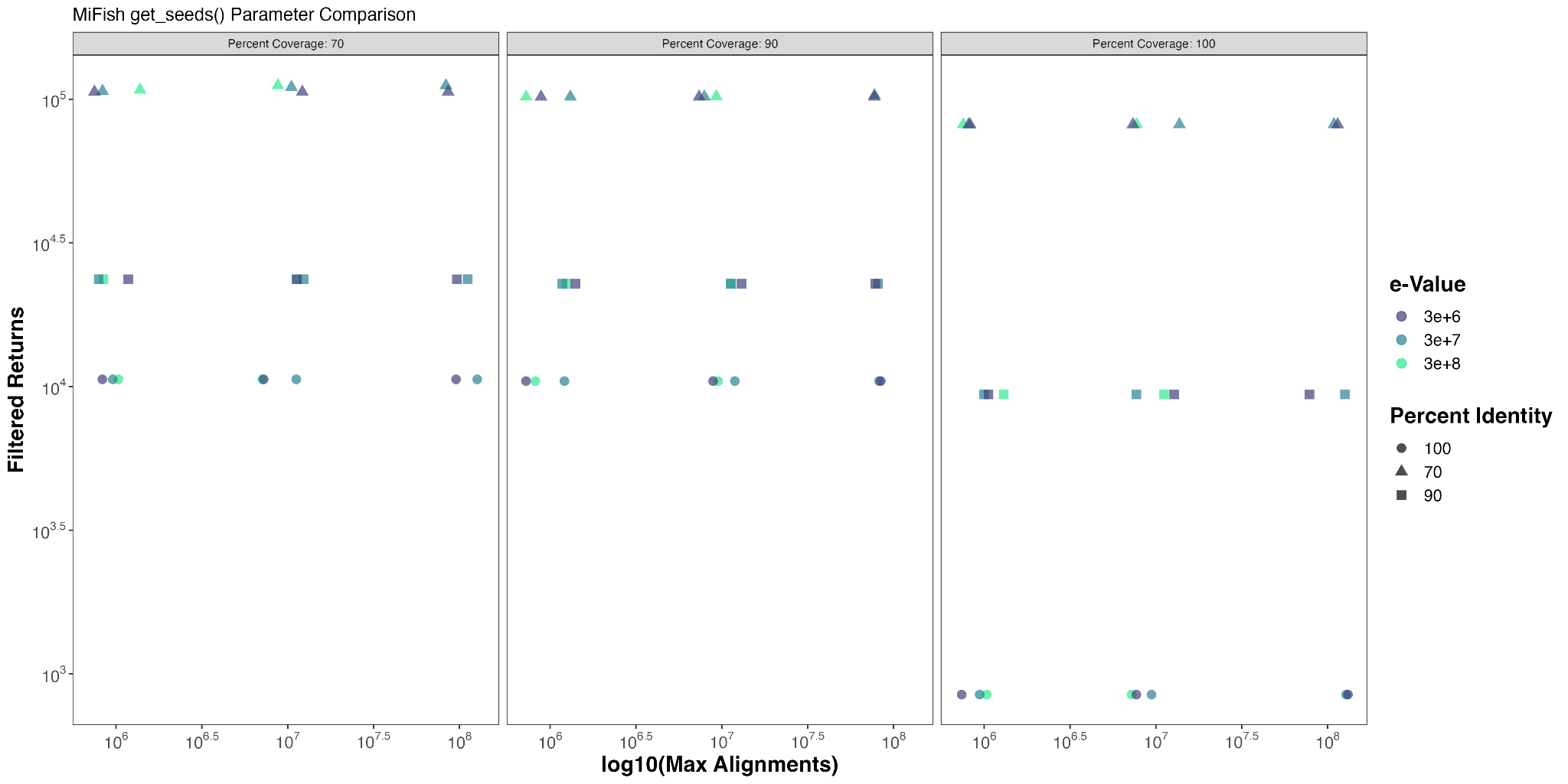

**Figure S1. rCRUX *get_seeds_local*() parameter effects on filtered returns for the MiFish primer set**

Comparison of number of filtered returns captured by rCRUX *get_seeds_local()* for a) MiFish 12S Universal Teleost locus. Percent cover and percent identity had a significant effect on the number of returned accessions (ANOVA, P <0.0001, Table S4). The max number of blast alignments and e-value had no significant effect on the number of returned accessions. We note that the greatest number of returned accessions occurred with an e-value of 3e+07 and max number of blast alignments of 10^8, percent identity of 70, and percent coverage of 70.

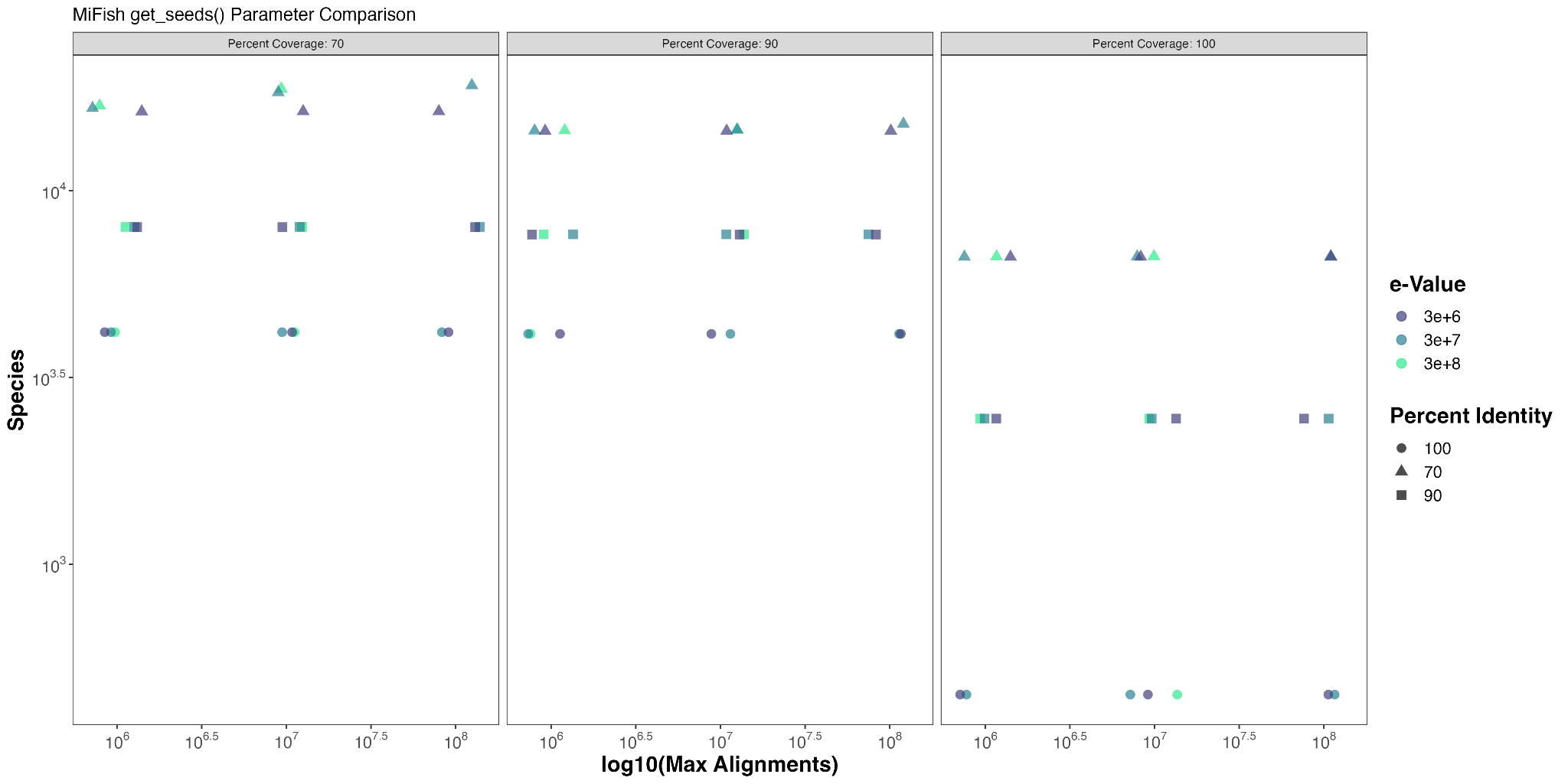

**Figure S2. rCRUX *get_seeds_local*() parameter effects on species for the MiFish primer set**

Comparison of number of species captured by rCRUX *get_seeds_local()* for a) MiFish 12S Universal Teleost locus. Percent cover and percent identity have the greatest effect on the number of returned species (ANOVA, P <0.0001, Table S4). The max number of blast alignments and e-value had no significant effect on the number of returned species. We note that the greatest number of returned species occurred with an e-value of 3e+07 and max number of blast alignments of 10^8, percent identity of 70, and percent coverage of 70.

Based on these results we chose to limit our comparisons of percent identity and coverage for *blast_seeds().* Here, we found a significant effect of the maximum number of blast alignments and taxonomic rank used for iterative blasting on the number of returns and species observed from *blast_seeds*() (ANOVA, P <0.0001, Figures S3 &S4, Table S5). We observed a significant effect of percent identity on the number of species observed (ANOVA, p<0.05), but not accessions (ANOVA, P>0.05). Four parameter sets shared the highest returned accessions and species. One of the four parameter sets was similar to the best performing *get_seeds_local()* parameter set with percent coverage of 70, percent identity of 70, e-value 3e+7, rank of genus, and max number of blast alignments = '100,000,000' (Table S5). We note that changing the max number of blast alignments = '10,000,000' produced nearly as many accessions and species as the above parameters, but was much less computationally intensive.

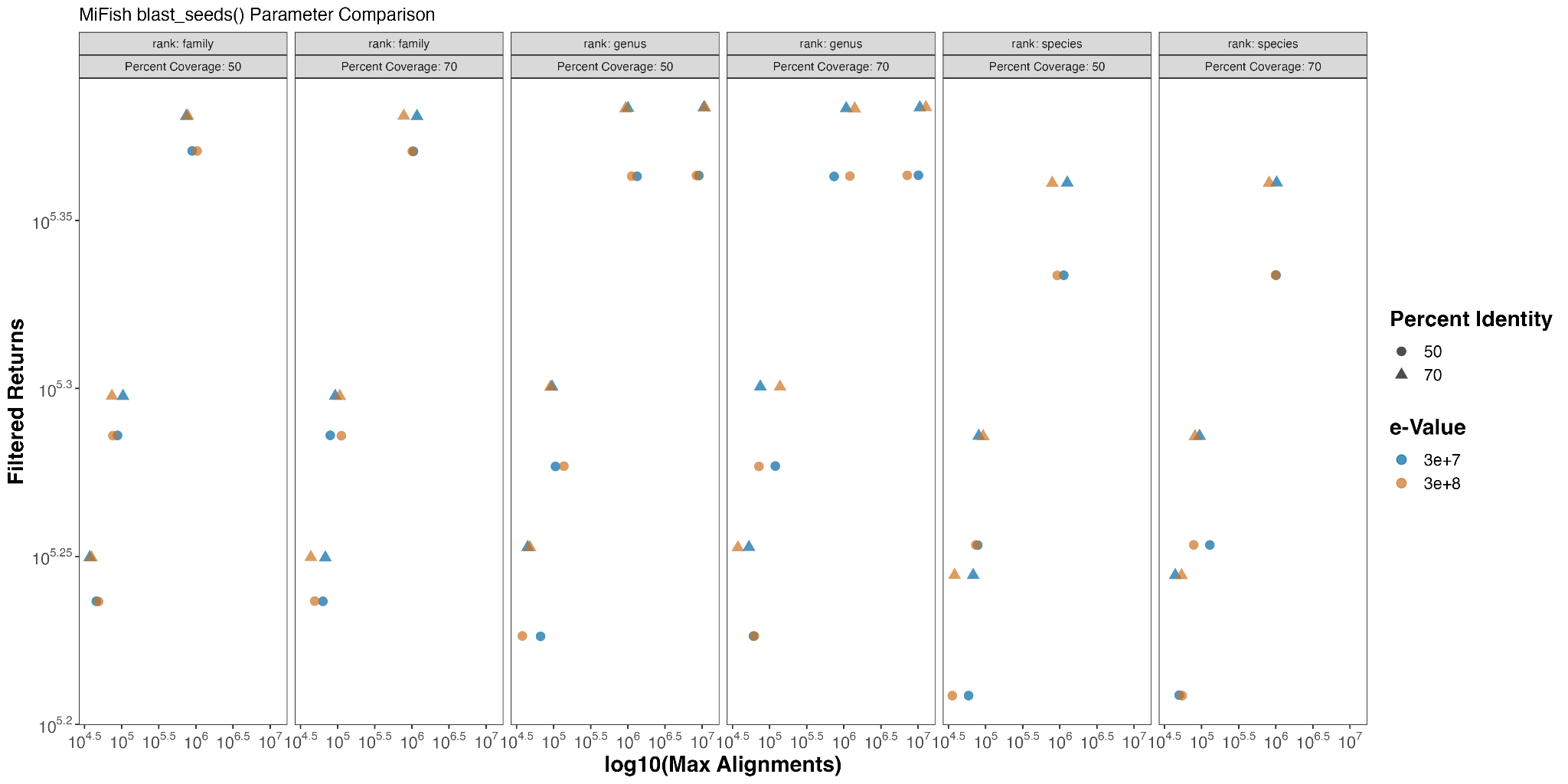

**Figure S3. rCRUX *blast_seeds*() parameter effects on filtered returns for the MiFish primer set**

Comparison of number of filtered returns captured by rCRUX *blast_seeds()* for a) MiFish 12S Universal Teleost locus. Number of alignments had the greatest effect on the number of observed returns (ANOVA, p <0.0001, Table S5). Taxonomic rank selected for iterative blasting also had a significant effect on the number of observed returns (ANOVA, p <0.05) with species having noticeably fewer returns than genus or family.

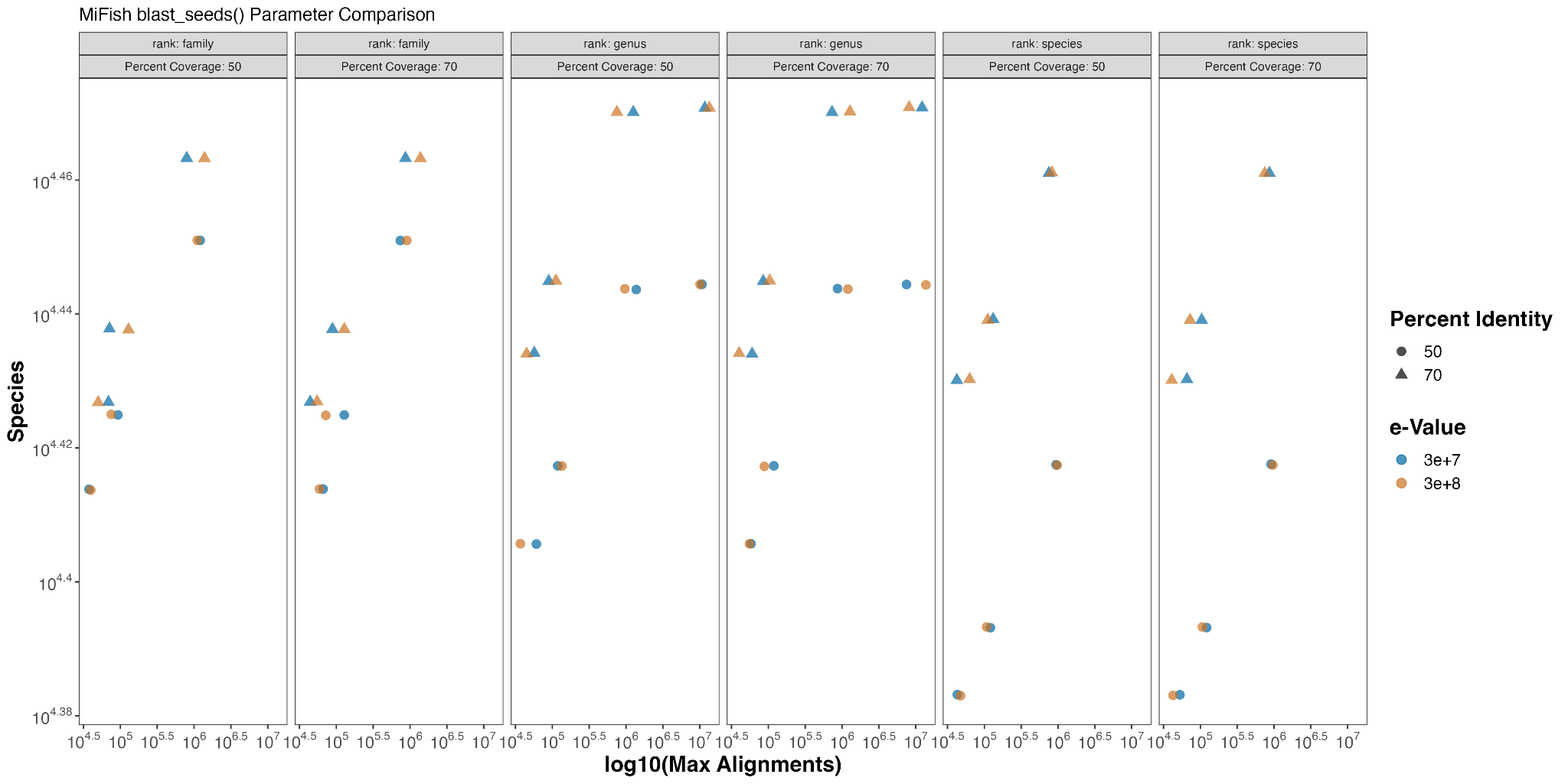

**Figure S4. rCRUX *blast_seeds*() parameter effects on species for the MiFish primer set**

Comparison of number of filtered returns captured by rCRUX *blast_seeds()* for a) MiFish 12S Universal Teleost locus. Number of alignments had the greatest effect on the number of observed returns (ANOVA, p <0.0001, Table S5). Taxonomic rank selected for iterative blasting also had a significant effect on the number of observed returns (ANOVA, p <0.05) with species having noticeably fewer returns than genus or family.

*trnl marker*

We found a significant effect of percent coverage and percent identity on the number of returned accessions and species observed from *get_seeds_local*() on the trnl locus (ANOVA, P <0.0001, Figures S5 &S6, Table S6). Percent cover, percent identity, and max number of blast alignments had a significant effect on the number of returned species and accessions (ANOVA, P <0.0001).

We observed the highest return accessions and species using percent coverage of 70, percent identity of 70, e-value 3e+8, and max number of blast alignments = '1,000,000'. The best performing parameters for the MiFish locus were the 2nd best performing for trnl (553 fewer species, and less than 2.5% total difference). We note that changing the max number of blast alignments to '10,000,000' and e-value 3e+6 produced nearly as many accessions and species as the above parameters, but was much less computationally intensive. As with the MiFish parameter testing, tests using parameters of e-value of 3e+8 timed out from a lack of computational resources more frequently than any other method (Table S4).

We found a significant effect of the percent identity and taxonomic rank used for iterative blasting on the number of returned accessions and species from *blast_seeds*() (ANOVA, P <0.0001, Figures S7 &S8, Table S7). We did not observe a significant effect of max alignments to blast on returned accessions or species (ANOVA, p<0.05), but found a significant effect on accessions (ANOVA, P>0.05). Four parameter sets shared the highest returned accessions and species (Table S7). One of the four top performing parameter sets utilized percent coverage of 50, percent identity of 70, e-value 3e+6, rank of species, and max number of blast alignments = '1,000,000'. Interestingly, percent identity of 50 and rank of species consistently resulted in greater returned accessions and species for the trnl locus, unlike the MiFish locus. The best performing parameter set from the MiFish locus to work on the trnl dataset (percent coverage of 70, percent identity of 70, e-value 3e+7, rank of genus, and max number of blast alignments = '10,000,000') was the 10th best performing parameter set (11,153 fewer species, and less than 14.9% total difference).

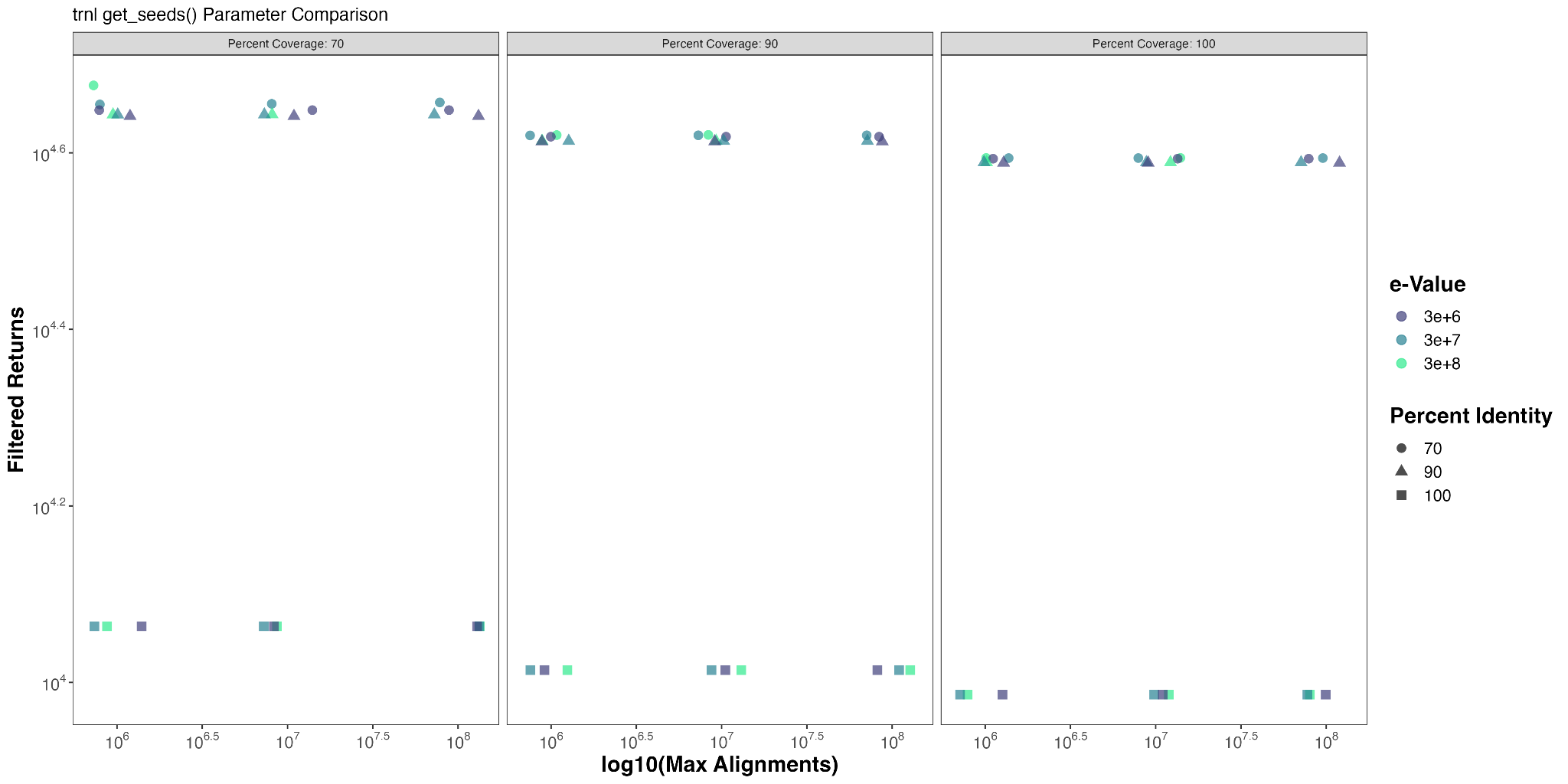

**Figure S5. rCRUX *get_seeds_local*() parameter effects on returned accessions for the trnl primer set**

Comparison of number of filtered returns captured by rCRUX *get_seeds_local()* for a) trnl locus. Percent cover, percent identity, and maximum alignments to blast had significant effects on the number of returned accessions (ANOVA, P <0.0001, Table S6). We observed the highest return accessions and species using percent coverage of 70, percent identity of 70, e-value 3e+8, and max number of blast alignments = '1,000,000'.

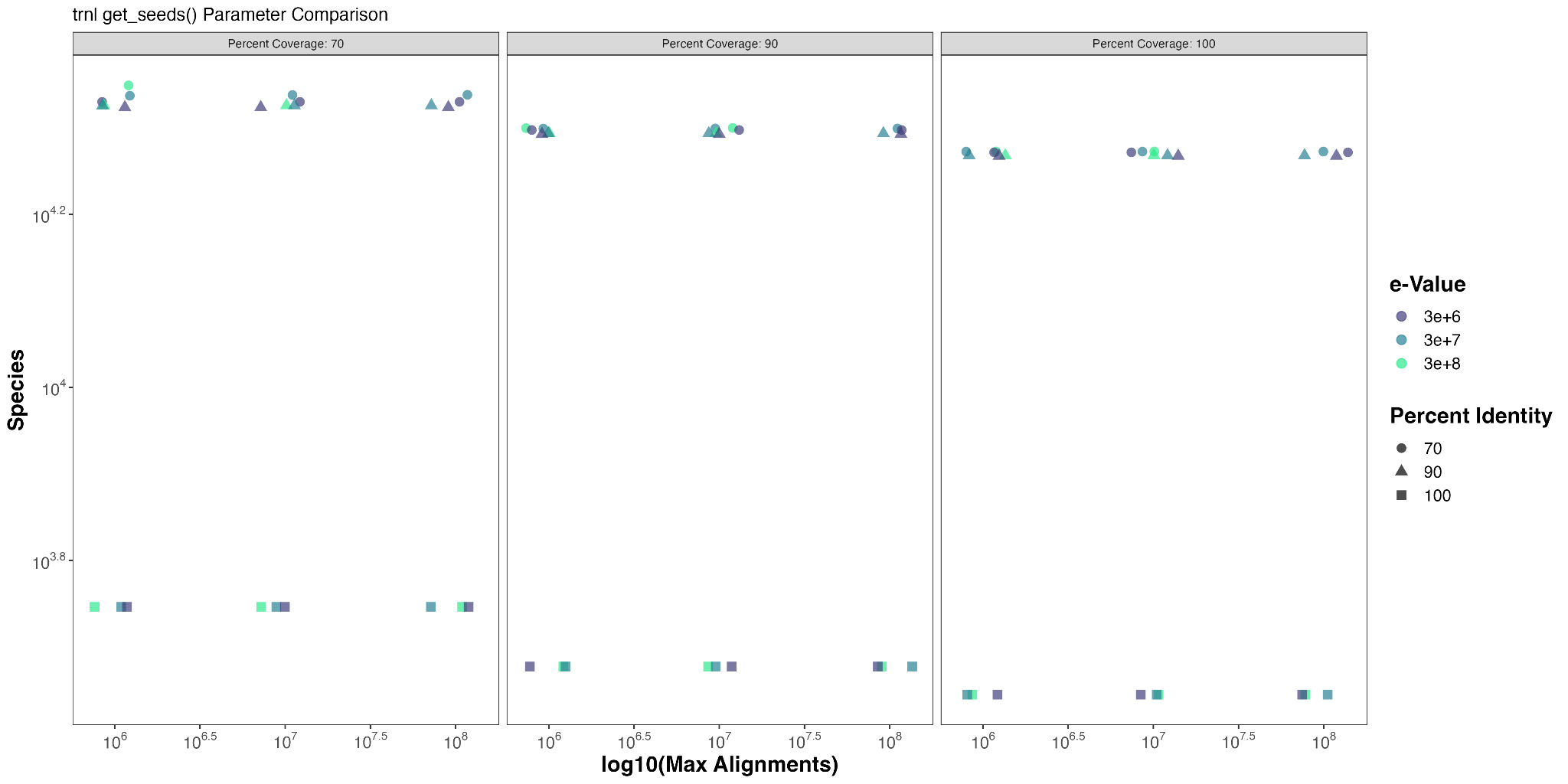

**Figure S6. rCRUX *get_seeds_local*() parameter effects on returned species for the trnl primer set**

Comparison of number of species captured by rCRUX *get_seeds_local()* for a) trnl locus. Percent cover, percent identity, and maximum alignments to blast had significant effects on the number of returned accessions (ANOVA, P <0.0001, Table S6). The highest return accessions and species using percent coverage of 70, percent identity of 70, e-value 3e+8, and max number of blast alignments = '1,000,000'.

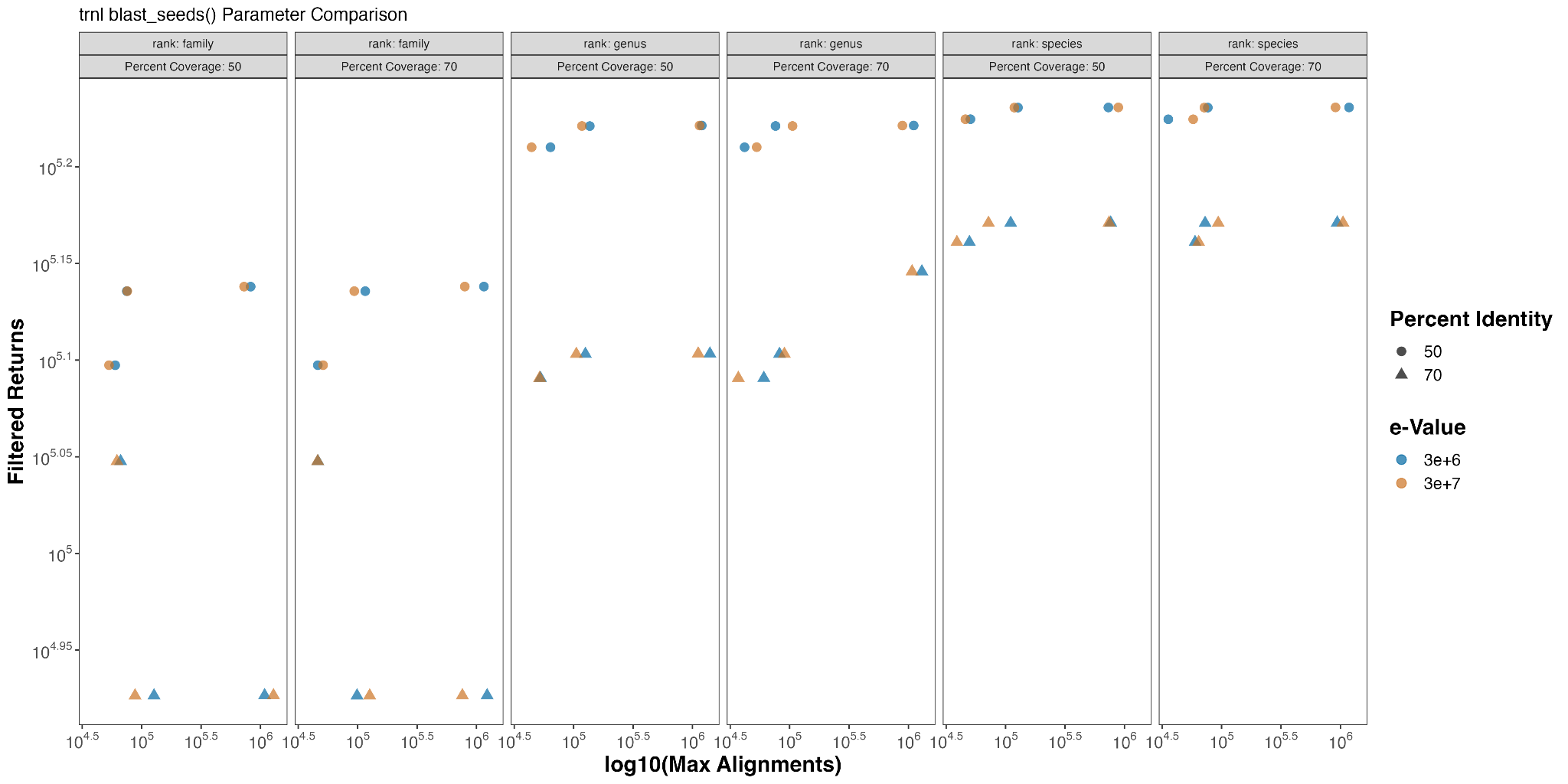

**Figure S7. rCRUX *blast_seeds*() parameter effects on returned accessions for the trnl primer set**

Comparison of number of returned accessions captured by rCRUX *blast_seeds()* for a) trnl locus. Percent identity and taxonomic rank for iterative blasting both had a significant effect on the number of returned accessions (ANOVA, p <0.0001, Table S7). Taxonomic rank had the greatest effect on returned accessions with rank of family having significantly fewer returned accessions than genus or species.

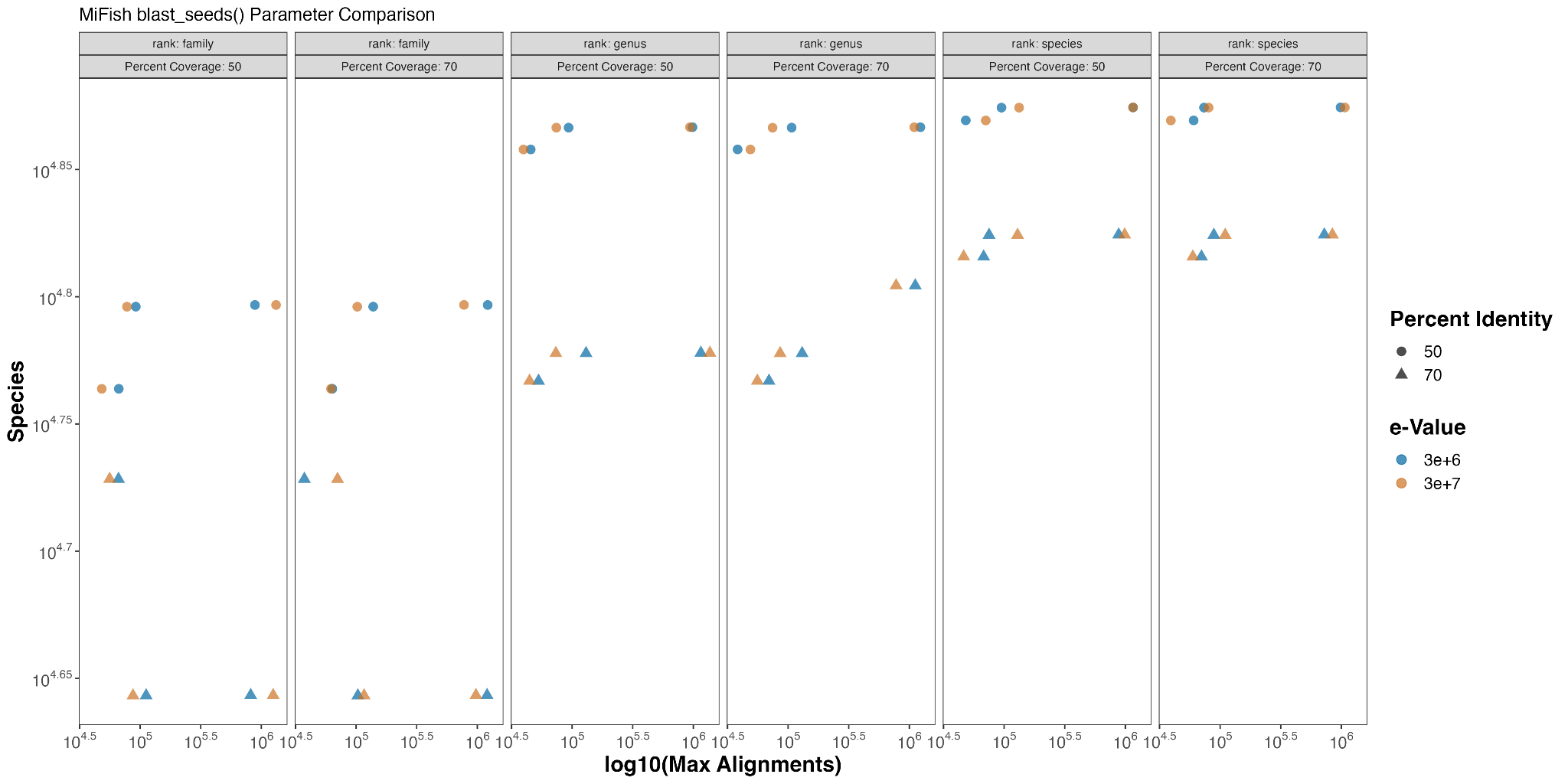

**Figure S8. rCRUX *blast_seeds*() parameter effects on species for the trnl primer set**

Comparison of number of returned species captured by rCRUX *blast_seeds*() for a) trnl locus. Percent identity and taxonomic rank for iterative blasting both had a significant effect on the number of returned species (ANOVA, p <0.0001, Table S7). Taxonomic rank had the greatest effect on returned species with rank of family having significantly fewer returned species than genus or species.

Given the similarity in optimized results from the parameter testing for both the MiFish and trnl locus for *get_seeds_local(),* we chose the following default parameters of percent coverage of 70, percent identity of 70, e-value 3e+6, and max number of blast alignments = '10,000,000'. We note that a lower e-value and max number of blast alignments were chosen to optimize efficiency between maximum number of returns and computational effort.

However, for *blast_seeds()* we had different best performing parameter sets. Interestingly, the trnl locus performed better with more permissive percent identity values which strongly suggests that there are more dissimilar trnl sequences than the 12S. At the same time, a greater number of returned accessions using iterative blasting at the rank of species suggests that there is greater diversity of trnl sequences within a given genus, resulting in fewer returns. Here, we opted to use the the following as default parameters: percent coverage of 70, percent identity of 70, evalue 3e+7, rank of genus, and max number of blast alignments = '10,000,000'. The lower e-value and max number of blast alignments were chosen to optimize efficiency between maximum number of returns and computational effort for the MiFish locus. Although this was not the best performing parameter set for trnl it still was in the top 10 returned accessions and filters and captured the vast majority of key taxa. These subtle differences in rCRUX performance pale in comparison to differences in rCRUX and other reference databases tested here (Figures 1-4).

These above defaults were used to generate all reference databases presented in the manuscript. However, we note that these results suggest that parameter tuning of both rCRUX functions result in a greater number of returned accessions and species and thus maybe warranted.

*Database Comparisons Filtered for Eukaryotes:*

The *get_seeds_local*() *in silico PCR* consistently captured a greater number of species than ECOPCR across the filtered MiFish 12S Universal Teleost, trnl, and FITS loci (Figure 1a,b,c). *get_seeds_local()* captured 95.3%, 88.2%, and 63.6% of species that ECOPCR captured while also capturing an additional 6,709 (40% of total species), 16,236 (74%), and 39,792 (56%) species respectively. Overlap in species captured ranged from 57% for the MiFish locus to 23% for the trnl locus (FITS was 28%).

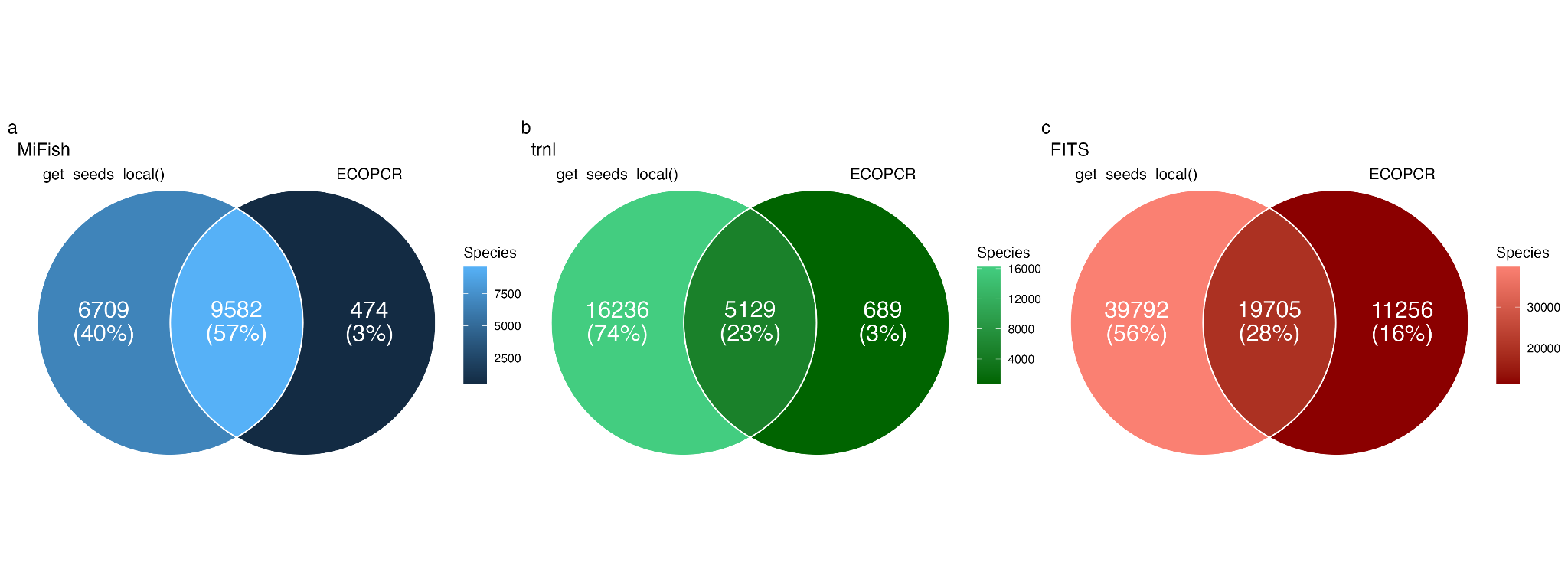

**Figure S9. rCRUX *get_seeds_local*() *in silico* PCR comparison with ECOPCR**

Comparison of number of species captured by rCRUX *get_seeds_local()* and OBITOOLS ECOPCR *in silico* PCR tools for a) MiFish 12S Universal Teleost, b) trnl, and c) fungal ITS (FITS) loci.

We further demonstrate that rCRUX reference databases consistently capture a greater number of accessions and species for MiFish (Figure 2), trnl (Figure 3), and FITS (Figure 4) loci across comparable benchmark reference databases.

For the MiFish reference database, only 32.5% of all species (n=7,933) shared across the 6 reference databases. Each reference database had unique sequences that were not shared with any other database (range: 12 - 3,452). rCRUX captured 86.6% (n=21,140) of all species observed across the MiFish reference databases. rCRUX uniquely had 14.2% (n=3,452) of all species observed and 6% (n=1,355) shared only with the CRUX reference database (Figure SXX).
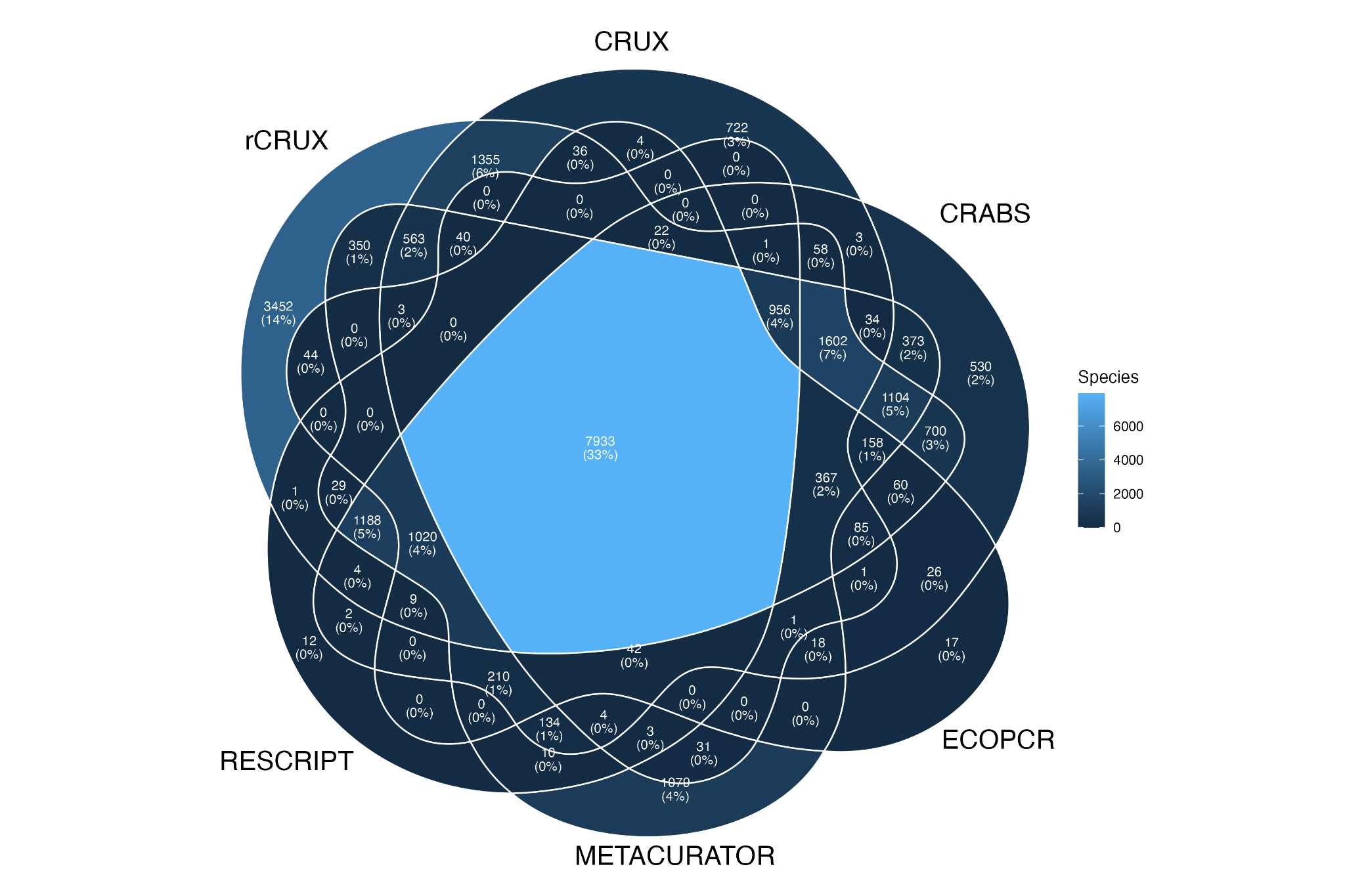

**Figure S10. MiFISH rCRUX *blast_seeds*() database comparison with CRUX, CRABS, ECOPCR, METACURATOR, and RESCRIPT databases.**

For the trnl reference database, only 133 species out of 67,872 species shared across the 5 trnl reference databases. Each reference database had unique sequences that were not shared with any other database (range: 55 - 42,575). rCRUX captured 93.9% (n=63,709) of all species observed across the trnl reference databases. rCRUX uniquely had 62.7% (n=42,575) species not observed in any other database and 18.7% (n=12,706) shared only with the METACURATOR database (Figure 3).

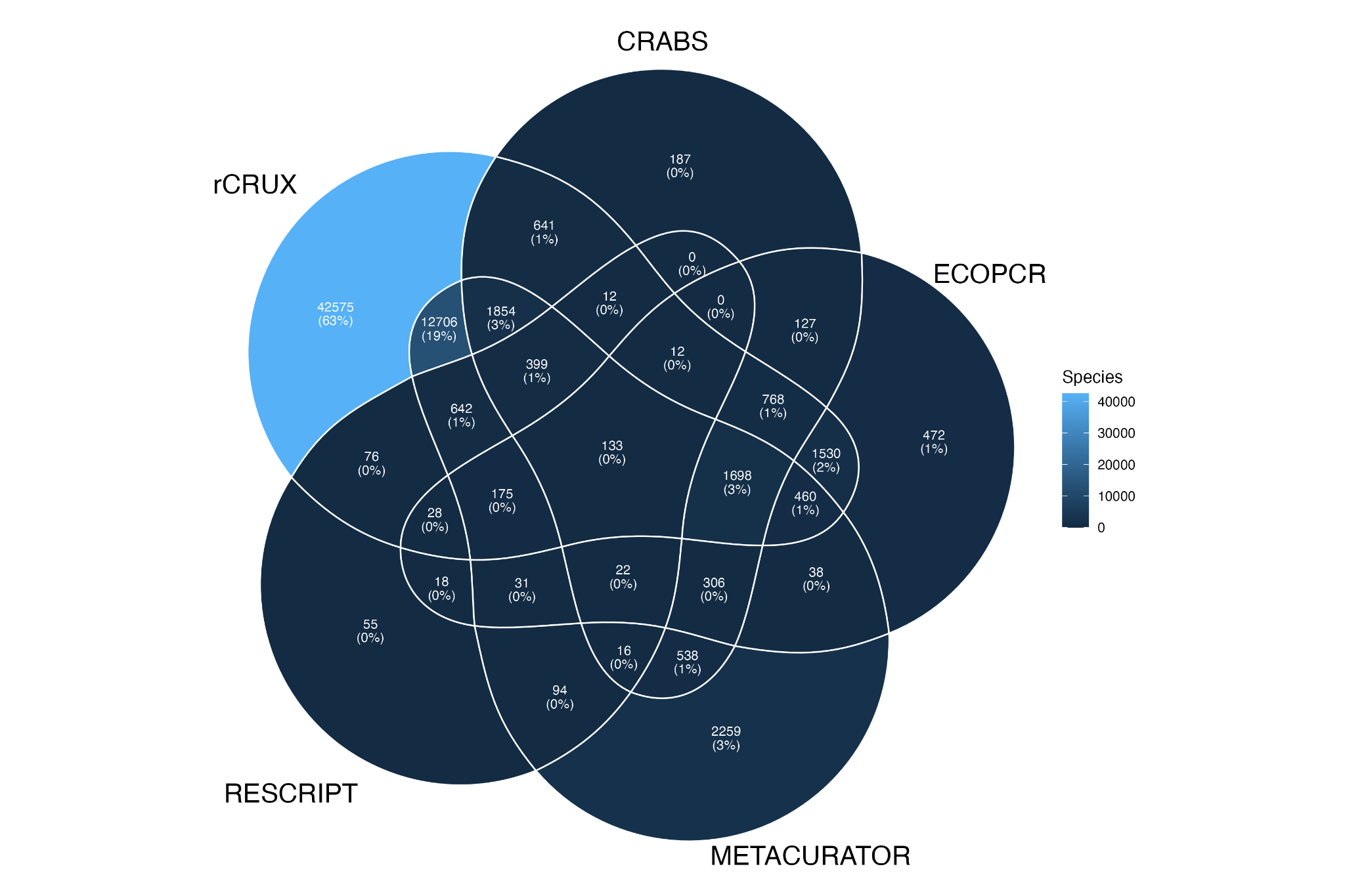

**Figure 11. trnl rCRUX *blast_seeds*() database comparison with CRABS, ECOPCR, METACURATOR, and RESCRIPT databases.**

For the FITS reference database, only 4.6% of all species (n=11,121) were shared across the 4 reference databases. Each reference database had unique sequences that were not shared with any other database (range: 920 - 177,272). rCRUX captured 94.6% (n=228,825) of all species observed across the FITS reference databases. rCRUX uniquely had 73.3% (n=177,272) species (Figure 3).

We note that the rCRUX databases were generated after the other databases, however they include the majority of species captured by compared methods. Together, these results benchmark rCRUX favorably against CRABS, METACURATOR, ECOPCR, RESCRIPT, and CRUX across a diversity of metabarcoding loci.

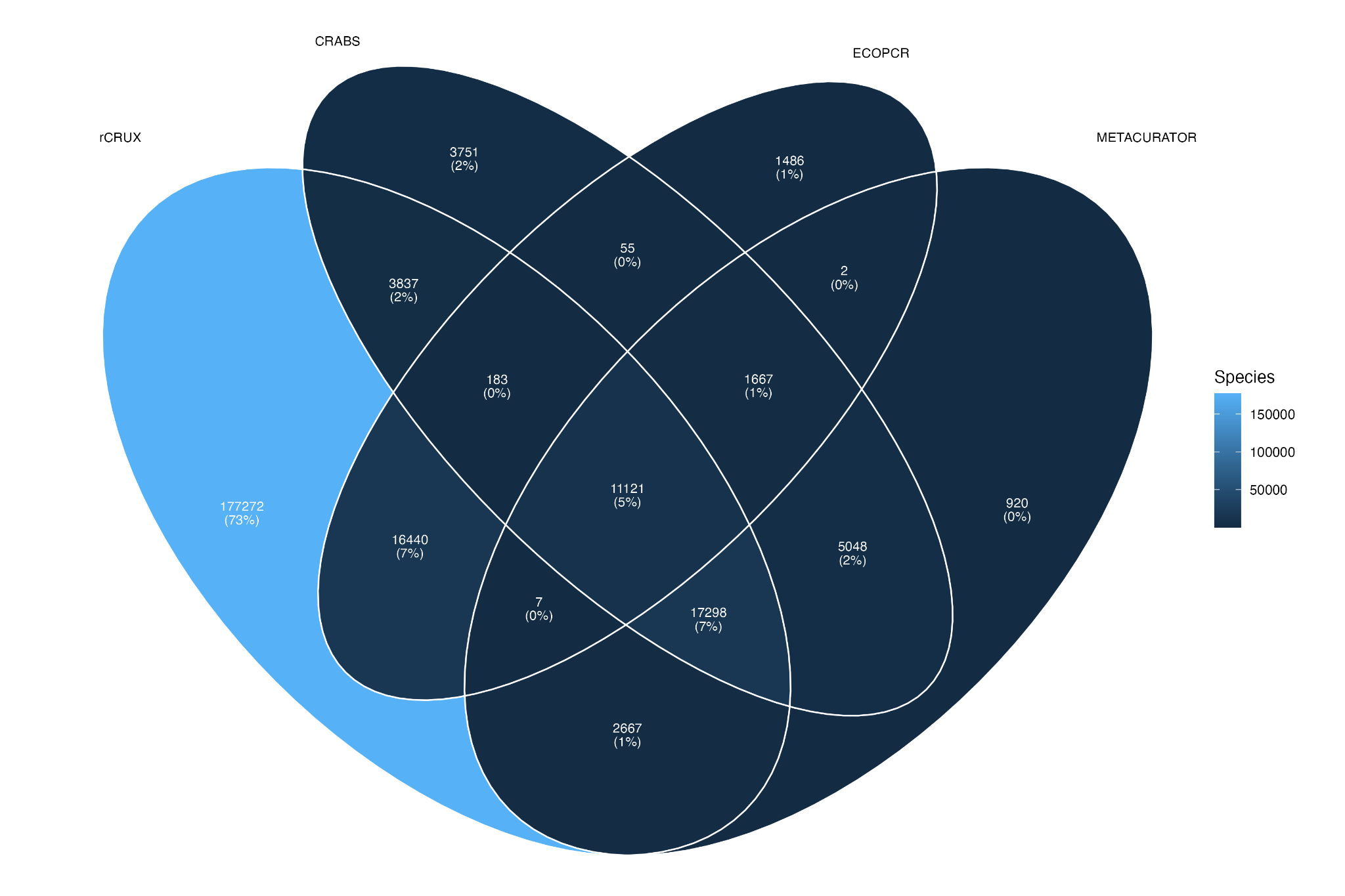

**Figure S12. FITS rCRUX *blast_seeds*() database comparison with CRUX, CRABS, ECOPCR, METACURATOR, and RESCRIPT databases**

Reproducibility:

We investigated the reproducibility of rCRUX database generation by running 10 replicate rounds of blast_seeds() for the MiFish Universal 12S marker (Supplementary Table X). Between replicates, the range of database returns was between 125,027-125,350 unique accessions averaging 125,171.7 returns with a coefficient of variation of 0.001. The number of species ranged between 21,126-21,236 with an average of 21,182.6 species with a coefficient of variation of 0.002. After running derep_and_clean() on the output the number of unique sequences ranged between 33,051-33,432 with an average of 33,301.70 and coefficient of variation of 0.004. The number of species ranged between 18,027-18,163 with an average of 18,109.7 species with a coefficient of variation of 0.003.

community composition in brood provisions of a small carpenter bee. Molecular Ecology 25:2302–2311.

Miya, M., Sato, Y., Fukunaga, T., Sado, T., Poulsen, J. Y., Sato, K., ... & Kondoh, M.

(2015). MiFish, a set of universal PCR primers for metabarcoding environmental DNA from fishes: detection of more than 230 subtropical marine species. *Royal Society open science*, 2(7), 150088.

R Core Team (2022). R: A language and environment for statistical computing. R

Foundation for Statistical Computing, Vienna, Austria.

URL https://www.R-project.org/.

Wickham H, François R, Henry L, Müller K, Vaughan D (2023). dplyr: A Grammar of

Data Manipulation. https://dplyr.tidyverse.org, https://github.com/tidyverse/dplyr
